# Cell-free gene regulatory network engineering with synthetic transcription factors

**DOI:** 10.1101/407999

**Authors:** Zoe Swank, Nadanai Laohakunakorn, Sebastian J. Maerkl

## Abstract

Gene regulatory networks are ubiquitous in nature and critical for bottom-up engineering of synthetic networks. Transcriptional repression is a fundamental function that can be tuned at the level of DNA, protein, and cooperative protein – protein interactions, necessitating high-throughput experimental approaches for in-depth characterization. Here we used a cell-free system in combination with a high-throughput microfluidic device to comprehensively study the different tuning mechanisms of a synthetic zinc-finger repressor library, whose affinity and cooperativity can be rationally engineered. The device is integrated into a comprehensive workflow that includes determination of transcription factor binding energy landscapes and mechanistic modeling, enabling us to generate a library of well-characterized synthetic transcription factors and corresponding promoters, which we then used to build gene regulatory networks *de novo*. The well-characterized synthetic parts and insights gained should be useful for rationally engineering gene regulatory networks and for studying the biophysics of transcriptional regulation.

## INTRODUCTION

Cell-free systems have emerged as versatile and efficient platforms for rapid engineering, characterization, and implementation of genetic networks. It has been demonstrated that linear genetic cascades (Noireaux et al. 2003), logic gates (Shin and Noireaux 2012), and oscillators (Karzbrun et al. 2014, Niederholtmeyer et al. 2013, 2015) could be implemented and characterized in cell-free systems, and that networks engineered in cell-free systems function in cells with remarkably similar characteristics, indicating that cell-free systems accurately emulate the cellular environment (Chappell et al. 2013, Niederholtmeyer et al. 2015). Besides these examples in molecular systems engineering and characterization of complex biological systems, cell-free systems provide a viable starting point for the bottom-up synthesis of artificial cells (Forster and Church 2006, Schwille et al. 2018). Work is progressing in establishing critical cellular sub-systems including DNA replication (van Nies et al. 2018), metabolism (Otrin et al. 2017), ribosome synthesis (Jewett et al. 2013), membrane synthesis (Bhattacharya et al. 2017), and protein structures (Furusato et al. 2018). Gene regulatory networks (GRNs) are one such critical sub-system, and here we demonstrate *de novo* bottom-up engineering and comprehensive characterization of synthetic GRNs in a cell-free system.

GRNs execute the genome and thus play a central role across all domains of life. Due to their importance and ubiquity, GRNs have been intensely studied and considerable progress is being made in deciphering components, topologies, and general mechanisms of GRNs, although a complete mechanistic understanding is still lacking. Because GRNs perform many sophisticated cellular tasks, synthetic biologists use GRNs to engineer new systems (Brophy and Voigt 2014) such as logic gates (Nielsen et al. 2016), toggle switches (Gardner et al. 2000), band-pass filters (Basu et al. 2005), and oscillators (Elowitz and Leibler 2000). Nonetheless, past and current efforts in engineering GRNs have shown that rational design is not yet possible, and that engineering GRNs still heavily relies on trial-and-error and high-throughput screening approaches (Nielsen et al. 2016). The inability to rationally design GRNs is in part due to the aforementioned lack of complete mechanistic understanding, and because basic GRN components such as transcriptional regulators and promoters are often neither fully characterized nor standardized. A corollary of the lack of an in-depth mechanistic understanding of these systems is that individual components are not yet readily composable. Nature provides a plethora of potential transcriptional regulators, but the number that have been tested and characterized remains rather limited. Most engineered GRNs make use of naturally occurring transcription factors, making it difficult to robustly engineer GRNs with such a non-standard set of proteins (Stanton et al. 2014). A library of well-characterized, synthetic transcription factors could alleviate many of these problems by providing a set of standardized transcription factors that are based on the same basic structural framework, and whose function can be extended by generating fusion proteins in a plug-and-play format.

Native GRNs employ a wide range of transcription factors that can be categorized into several structural families. The family with the largest number of members is the zinc-finger (ZF) family, followed by homeodomain, basic helix-loop-helix, and basic-leucine zipper (LZ) families (Vaquerizas et al. 2009). ZFs are of interest in biology as they represent the largest class of transcriptional regulators and are involved in diverse biological functions. ZFs are also appealing for bottom-up engineering as they consist of well-defined subunits that, in combination, determine DNA sequence specificity (Beerli and Barbas III 2002, Tebas et al. 2014). Many resources are therefore available that provide sequence specificity information for a large number of native (Najafabadi et al. 2015) and engineered (Fu and Voytas 2013) ZF transcription factors. An additional advantage is that ZFs are small (264 bp, 10.6 kDa (Zif268)) compared to other engineerable transcriptional regulators such as TALE (e.g. 1161–2397 bp, 39.9–82.6 kDa, DNA binding domain only (Moore et al. 2014)) or dCas9 (4107 bp, 158.3 kDa), so that the coding sequence for ZFs is easily obtainable and modifiable. Due to their small size and simple structure, ZFs can be readily expressed both *in vivo* and *in vitro*. Synthetic ZFs have already been successfully used as activators in *S. cerevisiae* (Khalil et al. 2012) and human cells (Lohmueller et al. 2012). Here we engineer and explore the use of synthetic ZF transcriptional regulators as ideal building blocks for bottom-up design and implementation of cell-free GRNs.

In this paper, we took advantage of an existing synthetic ZF library (Blackburn et al. 2015) to generate a well-characterized resource of transcriptional repressors and corresponding synthetic promoters that can be used for bottom-up design, implementation, and characterization of GRNs in cell-free systems. While the mechanism of action of the simplest prokaryotic repression is competitive inhibition (Ptashne et al. 1976), it has long been appreciated that both *cis* modifications to the promoter, such as operator position (Cox III et al. 2007), basal promoter strength (Lutz and Bujard 1997), as well as *trans* modifications to the transcription factor itself strongly affect repression (Lanzer and Bujard 1988, Sharon et al. 2012). These inter-dependencies result in a large experimental space with many degrees of freedom. In order to tackle this complexity we developed a microfluidics based method capable of performing 768 cell-free transcription-translation (TX-TL) reactions on a single device. The ability to rapidly generate ZF repressor and promoter variants using fast PCR assembly and the use of our high-throughput microfluidic device allowed us to perform a comprehensive characterization of repressors and promoters. We investigated the effects of binding site position, binding site affinity, binding site combinations, and cooperative interactions between the repressors on transcriptional repression performance. We generated quantitative position weight matrices (PWMs) for four ZF repressors with MIT-OMI (Maerkl and Quake 2007), which allowed us to rationally tune binding site affinity and promoter output. Finally, we used the parts library and insights acquired in this study to engineer logic gates, showing that *de novo* synthetic GRNs can be rationally engineered using a bottom-up approach. The transcription factor / promoter parts library, data, and methods described here provide a resource that should facilitate efforts to build synthetic GRNs, serve as a viable approach for building GRNs for use in artificial cells, and establish an experimental platform for studying the biophysics of transcriptional regulation.

## RESULTS

### A. Design and characterization of a microfluidic device for high-throughput cell-free experiments

The design space of even a single TF – promoter pair is large, encompassing different binding site affinities, binding site positions, binding site sequences, and binding site combinations. This complexity necessitates high-throughput methods capable of the functional characterization of hundreds to thousands of engineered variants. Current approaches in cell-free synthetic biology primarily rely on standard microtiter plates, which require a minimal reaction volume of 5 – 10 μL. Such relatively large volumes quickly become cost-limiting in terms of how much cell-free reaction solution and DNA is required to perform the assays. Researchers recently made use of an acoustic liquid handling robot that reduced reaction volumes to 2 *μL* in a 384 well plate format (Moore et al. 2018). Here we repurposed the MITOMI platform, a microfluidic device originally developed for high-throughput molecular interaction analysis (Garcia-Cordero and Maerkl 2016, Maerkl and Quake 2007), and applied it to the high-throughput characterization of cell-free genetic networks. The repurposed device performs 768 cell-free reactions, and reduces volumes by ∼4 orders of magnitude to ∼690 pL per reaction.

The process involves the synthesis of DNA parts, followed by microarraying and incorporation into microfluidic unit cells where they serve as templates in cell-free TX-TL reactions (Figure 1A). To expedite the synthesis of large libraries of DNA parts we used an assembly PCR strategy to generate linear DNA templates with different promoter regions upstream of a deGFP gene. A microarray robot is used to spot the linear templates onto an epoxy-coated glass slide, on top of which the PDMS device is aligned. Immobilizing DNA within each reaction chamber first requires surface patterning in the assay section of each unit cell, resulting in a circular area of neutravidin to which biotinlyated DNA can bind. Once DNA is surface immobilized, cell-free extract is flowed into the device and the unit cells are isolated from one another while the TX-TL reactions occur. A detailed schematic of the experimental procedure is shown in Figure S1A.

**Figure 1:**
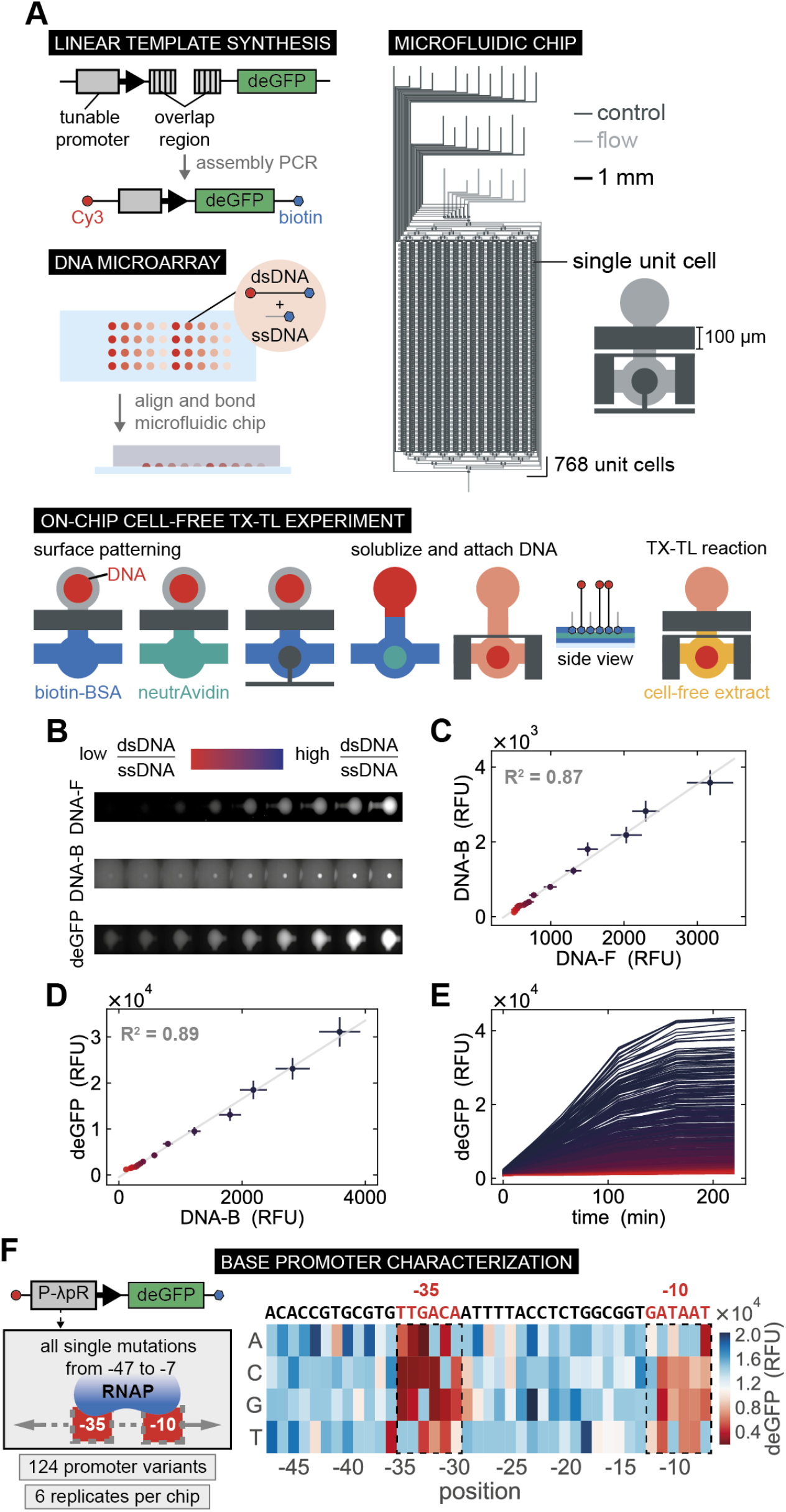
High-throughput microfluidic cell-free reactions. (**A**) A schematic overview of the experimental design including synthesis of DNA templates, DNA spotting, chip alignment, surface functionalization followed by DNA immobilization and on-chip cell-free TX-TL reactions. (**B**) Fluorescence images of *Cy3-DNA_F_*, Cy3-DNA_B_, and deGFP expressed for a range of dsDNA:ssDNA ratios. (**C**) Quantification of surface immobilized DNA (*DNA_B_*) as a function of free DNA in solution (*DNA_F_*). (**D**) deGFP expression at the final time point as a function of *DNA_B_* concentration. All values represent means ± SD (*n* = 48). (**E**) deGFP expression measured over time in all unit cells. (**F**) Schematic of a promoter library design and on-chip experimental throughput, followed by the deGFP expression for all single base mutations from position −47 to −7 of the *λP_R_* promoter.

Controlling the precise amount of DNA in each unit cell is important for quantitative experiments. By simply varying the concentration of spotted biotinylated DNA templates we were unable to precisely control DNA concentration on-chip. We thus developed an approach based on spotting a mixture of single stranded biotinylated DNA oligos (ssDNA) and double stranded DNA templates (dsDNA). The amount of DNA immobilized on the surface reached saturation at a concentration of ∼100 nM spotted DNA (Figure S2). We therefore held the total concentration of spotted DNA above this saturation point. Changing the ratio of dsDNA:ssDNA gave rise to a linear correlation between the concentration of dsDNA free in solution (*DNA_F_*) and dsDNA bound to the surface (*DNA_B_*), and was insensitive to the total amount of DNA deposited during spotting (Figure 1B, C). This approach allowed us to immobilize DNA over a wide concentration range, which gave rise to corresponding levels of expressed deGFP (Fig. 1D, E). The results obtained with the high-throughput microfluidic device are reproducible with a global normalized root-mean-square deviation of ∼14%, not only when a single dsDNA template is used, but also for more complex experiments requiring multiple templates in each unit cell (Figure S3). Furthermore, a subset of on-chip measurements was carried out in standard micro-well plate reactions, showing good correlation (Figure S4).

To demonstrate the high-throughput capabilities of our microfluidic chip we created and characterized a library based on the *E. coli σ*_70_ *λP_R_* promoter. We synthesized 124 promoter variants that covered all possible single base mutations within the −47 to −7 region of the *λP_R_* promoter (Figure 1F). Cell-free reactions for each promoter were run in 6 replicates on a single chip and yielded deGFP expression profiles revealing the impact of each mutation on protein expression (Figures 1F, S3A). As expected, mutations within the −10 and −35 boxes affected deGFP expression most strongly and the results are comparable to previous results obtained by an in vivo analysis of the lac promoter (Kinney et al. 2010).

Protein synthesis eventually stops in cell-free batch reactions as seen in the saturation dynamics in time course measurements (Figure 1E); this is fundamentally different from cellular steady state protein levels which result from balancing production with degradation and dilution rates. In this paper we report end-point batch reaction values and derived quantities such as fold repression. It is thus important that the end-point values correspond to protein production rates. While the relationship between the initial rate of deGFP production and its final saturated level may be complex, we observe a linear relationship between the two quantities under our experimental conditions (Figure S5). This is an important validation of our use of end-point protein levels and linearly derived quantities such as fold repression as proxies for synthesis rates and their ratios.

### B. Zinc-finger repressor and promoter library design

Using the characterization of the *λP_R_* promoter as a starting point, we applied our chip to the in depth characterization of synthetic ZFs for use as transcriptional repressors. We adopted a ZF design based on Zif268, a three-finger Cys2His2 protein. A large ZF repressor library can be generated by combinatorially shuffling a small number of individual ZF domains (Figure S6A). We utilized ZF proteins drawn from a 64-member library that we previously synthesized and characterized (Blackburn et al. 2015) (Figure S6).

The affinity of a ZF repressor to DNA can be improved by increasing the number of finger domains (Kamiuchi et al. 1998, Kim and Pabo 1998, Moore et al. 2001, Pomerantz et al. 1998). The same effect can also be achieved by engineering dimerizing ZFs that bind cooperatively. An early example used structure-based design to engineer a two-finger ZF which dimerized via a LZ motif to form a four-finger complex (Wolfe et al. 2003, 2000). Three-finger ZFs have also been dimerized using PDZ domains (Khalil et al. 2012). Cooperative interactions are of interest because they potentially increase the nonlinearity of regulation, as well as decreasing non-specific binding compared to extended arrays of ZFs. To study cooperative interactions we built several different ZFs fused to either PDZ or LZ domains (Figure S6B).

In parallel, we designed corresponding repressible promoter libraries. As we use an *E. coli* cell-free system (Sun et al. 2013), we based our promoter designs on the strong λP_R_ promoter in combination with transcription and translation elements optimized for *E. coli* cell-free expression (Sun et al. 2014). Previous work has shown that the most effective position for transcriptional repression is the space between the −35 and −10 boxes (Cox III et al. 2007); we thus generated a library with consensus ZF binding sites (ZFBSs) inserted into this location. Additionally, we built promoters with a second ZFBS upstream of the −35 box, allowing us to study the effect of multiple non-cooperative and cooperative ZFBSs (Figure S6B). The promoters drive expression of a deGFP reporter, a GFP protein previously optimized for cell-free translation (Shin and Noireaux 2010). All constructs were built and tested using linear DNA templates generated by PCR in concordance with recommended guidelines for cell-free expression (Sun et al. 2014).

### C. Repression with single and multiple binding sites

We performed an in depth characterization of 11 synthetic ZFs by assessing their repressive capacity in cell-free reactions, and by measuring their respective dissociation constants (*K_d_*) with MITOMI. We used MITOMI to measure the *K_d_s* for each ZF against all possible target promoters. By localizing pre-synthesized his-tagged ZFs to the surface of each unit cell we are able to measure the binding of DNA sequences spanning the promoter region including the ZF binding site (Figure 3A, S1B). We obtained standard Gibbs free energies, Δ*G* = *RT* ln(*K_d_*), for each ZF - target promoter complex (Figure 3B). A range of binding strengths was observed for the respective consensus ZF binding sequences, as well as low affinity off-target binding. The CBD zinc finger was included as a negative control as it does not bind to its own predicted binding site nor any of the other targets.

**Figure 2:**
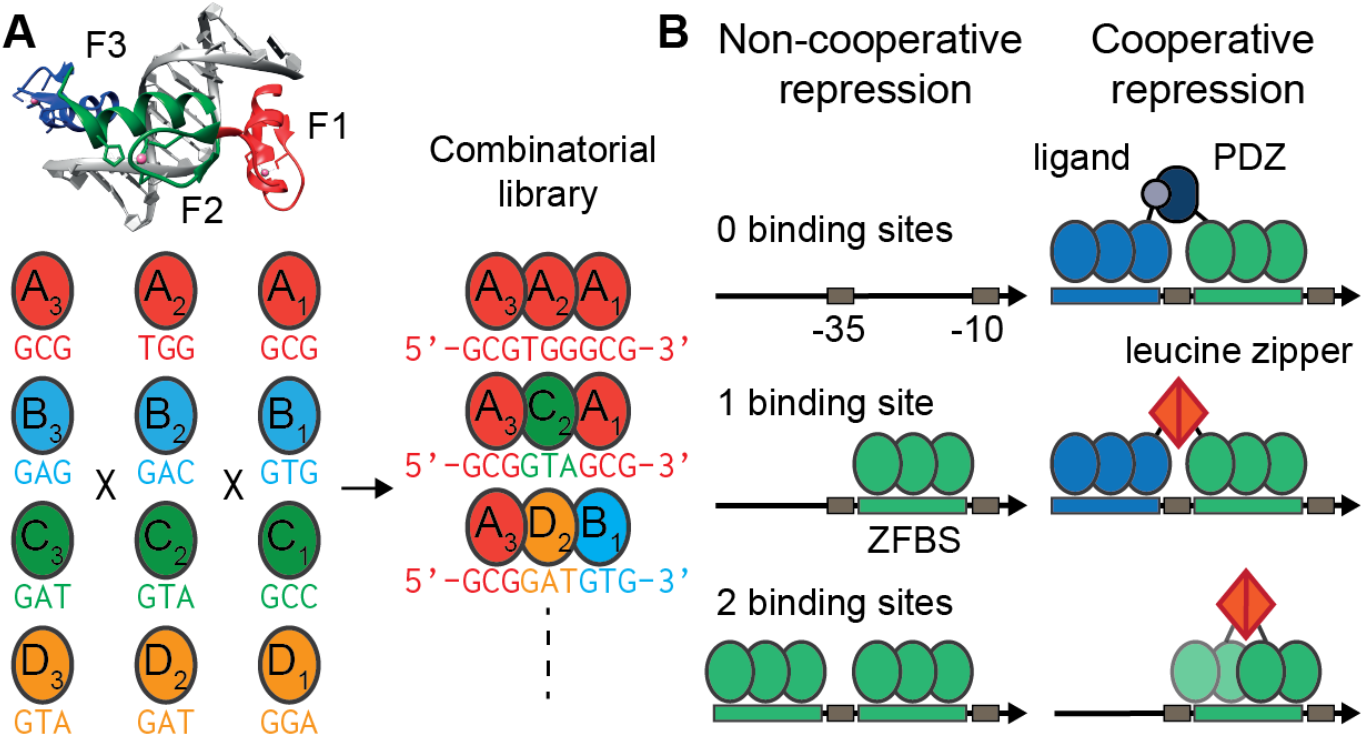
Zinc-finger repressor and promoter design. (**A**) Our repressor design is based on the Zif268 protein, which contains three Zn-fingers (`F1–F3`), each recognizing a nucleotide triplet. We used a combinatorial library of repressors by starting from four initial ZF proteins (here labelled with the codes `AAA–DDD`), and shuffling the individual Zn-fingers while preserving their position within the protein. (**B**) We designed a library of repressible promoters based on the *λP_R_* promoter. To test the effectiveness of repression we designed promoters containing single and dual sites with variable spacing, as well as engineering direct cooperativity between ZF proteins, which can be mediated by PDZ-ligand or LZ interactions.

**Figure 3:**
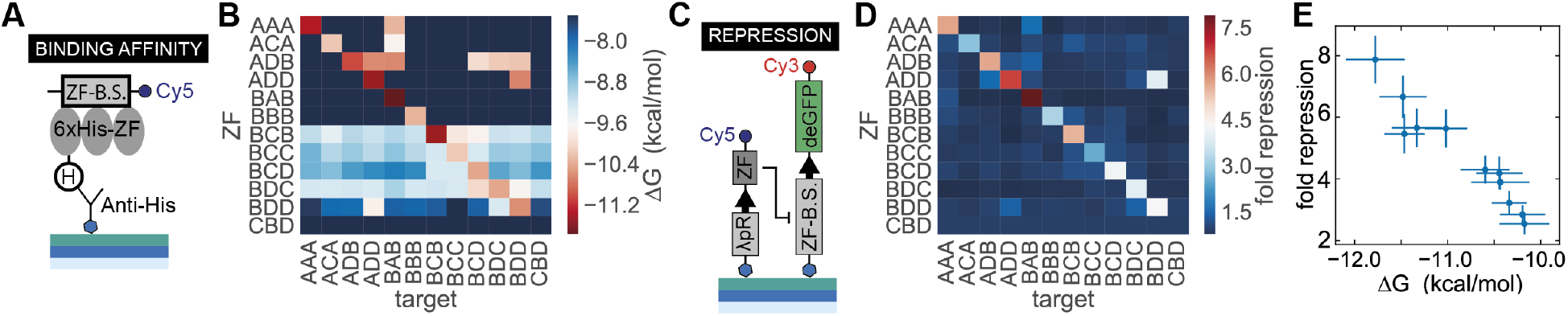
Zn-finger binding affinities, functional repression and orthogonality. (**A**) Schematic depicting the MITOMI assay used to determine TF - DNA binding affinities. (**B**) Affinity orthogonality matrix of Δ*G* values for all zinc fingers versus all possible DNA targets. (**C**) Schematic depicting the linear templates used to test functional repression in on-chip cell-free TX-TL reactions. (**D**) Fold repression orthogonality matrix for all zinc fingers versus all possible targets. (**E**) Fold repression values versus measured *K_d_* for all ZF - promoter consensus pairs. The fold repression data was collected from a single chip and all values represent means ± SD (*n* = 5). The error bars shown for the *K_d_* values represent the 95% confidence interval for the fit to a single binding site model.

To test whether the relative binding strength of each ZF related to functional gene repression, we implemented cell-free TX-TL reactions screening the same matrix of ZFs versus promoters. Each microfluidic unit cell contained a linear template encoding the ZF to be tested and a second linear template encoding deGFP downstream of a promoter with a single ZF binding site (Figure 3C). Binding of the expressed ZF to the target promoter would lead to down-regulation of deGFP expression. A common measure of repression performance is fold repression, or the ratio of unrepressed to repressed expression levels. Unrepressed measurements were obtained by co-expressing the target promoter template with the nonbinding *ZF_CBD_* template to control for loading effects (Siegal-Gaskins et al. 2014). Despite some off-target binding observed by MITOMI, functional repression of all ZF – target pairs was almost perfectly orthogonal (Figure 3D), with one exception: the repression of promoter *BDD* by *ZF_ADD_*. However the general trend of weak off-target affinities translated to no or minimal off-target repression, resulting in functional repression only for cognate pairs. Furthermore, on-target fold repression directly correlated with the measured MITOMI affinity values (Figure 3E). Using two high-throughput microfluidic techniques we were able to characterize the binding affinity, repressive strength, and orthogonality of synthetic transcription factor – promoter pairs.

Promoters with a single ZF binding site achieved low to medium fold repression levels in the range of 1.5 to 7 (Fig. 4A). We tested whether placing an additional binding site upstream of the −35 box could further improve fold repression levels. While fold repression is a convenient measure used to describe the functionality of a given repressor - promoter pair, for applying these repressors in genetic networks it is important to also consider basal promoter strength (unrepressed state) and leak (repressed state). These quantities are also shown in Figure 4, where we observed that variation in binding site sequence led to variations in basal promoter strength; this variation increased upon inclusion of the second binding site upstream of the −35 box. At the same time, the average leak from the repressed state decreased for the dual site library, resulting in higher fold repression values. Overall, fold repression improved for almost all two binding-site promoters, with the best promoters achieving a fold repression level of 7 – 10 (Figure 4B). These results showed that good repression levels can be achieved by synthetic ZF repressors with either single or double binding site promoters in a cell-free system.

**Figure 4:**
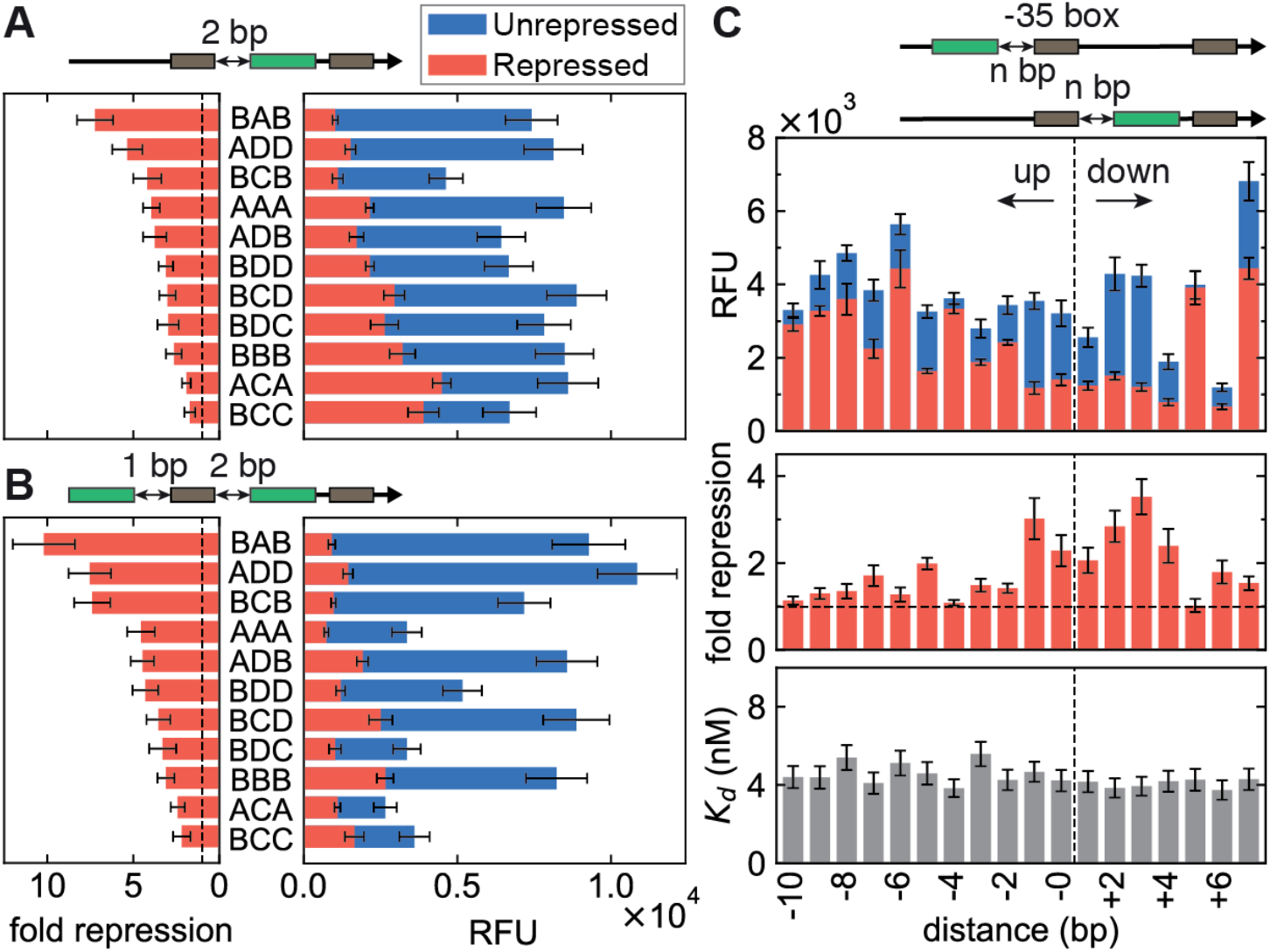
Effect of binding site number and position on repression. Shown are endpoint unrepressed and repressed levels for the single binding site library (**A**), and the dual site library (**B**). The data were rank-ordered by fold repression values within each library. Data were collected from two separate chips and all values represent means ± SD (n = 10). (**C**) A single `BCB` site was shifted up to 10 bp upstream and 7 bp downstream of the −35 box `TTGACA`; position values are given by the number of nucleotides separating the 9-bp site from the −35 box. The binding affinities of the ZF to its target site remains approximately constant irrespective of target site position ((**C**), bottom). Data from the top panel was measured from a single chip and all values represent means ± SD (*n* = 7). The error bars shown for the *K_d_* values represent the 95% confidence interval of the fit to a single binding site model.

Next we characterized the effect of binding site position on repression strength. We generated a library of promoters containing a single ZF binding site that was placed in various positions relative to the −35 box. Best fold repression was achieved by positioning binding sites directly proximal to the −35 box, in the range of −2 to +4 bps relative to the start and end of the −35 box, respectively. We also observe that repression is sensitive to single bp shifts in position. For instance, the site at the +5 position is effectively non-functional compared to repressing neighbouring sites at +4 and +6; and the site at the −5 position exhibited significantly stronger repression than its neighbours at −4 and −6. Based on the crystal structure alignment of ZF and RNA polymerase bound to DNA containing the binding site at position +5, we note that it is possible for both proteins to bind simultaneously with minimal steric interference. To ascertain that the observed repression strengths were not due to changes in binding site affinity of the ZF, as each binding site is located in a different sequence context, we measured the binding affinity of the ZF repressor to each promoter using MITOMI. The results showed only minor differences in affinity across all promoters, suggesting that the ZF repressor bound to these promoters with equal strength. Promoter repression thus appears to be primarily a function of the ability of the ZF to sterically hinder and compete with RNA polymerase. These data are consistent with an occlusion mechanism whereby RNAP binding is competitively inhibited by ZF binding (Ptashne et al. 1976), and the effectiveness of the competition is dependent on the relative positions of ZF and RNAP on the promoter.

### D. Engineering cooperativity

We showed that incorporating a second binding site can result in improved fold repression. However, engineering certain types of genetic circuits often requires an additional increase in the nonlinear response as well as a decrease in the leak for a given promoter – TF pair. Nonlinearity can be increased by introducing cooperativity via protein – protein interactions. We implemented two different protein interaction domains previously demonstrated to successfully dimerize ZFs.

PDZ domains enable natural protein – protein interactions by binding specific C-terminal peptide sequences with micromolar affinity (Khalil et al. 2012). We took advantage of this interaction to engineer cooperativity by linking *ZF_BCB_* to a mammalian *α*1-syntrophin PDZ domain, and *ZF_ADD_* to its corresponding cognate C-terminal peptide ligand (VKESLV). Furthermore, we linked *ZF_ADD_* with a non-cognate ligand (VKEAAA) to use as a noncooperative control.

The second type of interaction we explored was dimerization by linking *ZF_BCB_* and *ZF_ADD_* to GCN4 LZ domains. The GCN4 LZ has previously been used in a structure-based design to enable homodimerization of two-finger ZFs (Wolfe et al. 2000), and we thus also tested this existing structure. In both cases, a mutated LZ was used as a negative control.

Preliminary studies on a plate reader demonstrated that ZFs containing interaction domains exhibited significantly increased fold repression and decreased leak (Figure 5A, B). Whereas two non-cooperative repressors gave a maximum fold repression of ∼6, this value was increased to ∼30 for PDZ and ∼16 for LZ-mediated cooperativity. Concurrently, leak values decreased four-fold from around 4000 to <1000 RFUs. One critical parameter affecting PDZ cooperativity was the choice of linker, with an optimized glycine-serine linker vastly outperforming a rigid proline linker. The two-finger LZ transcriptional repressor also performed very well, achieving a fold repression ratio of ∼28.

**Figure 5:**
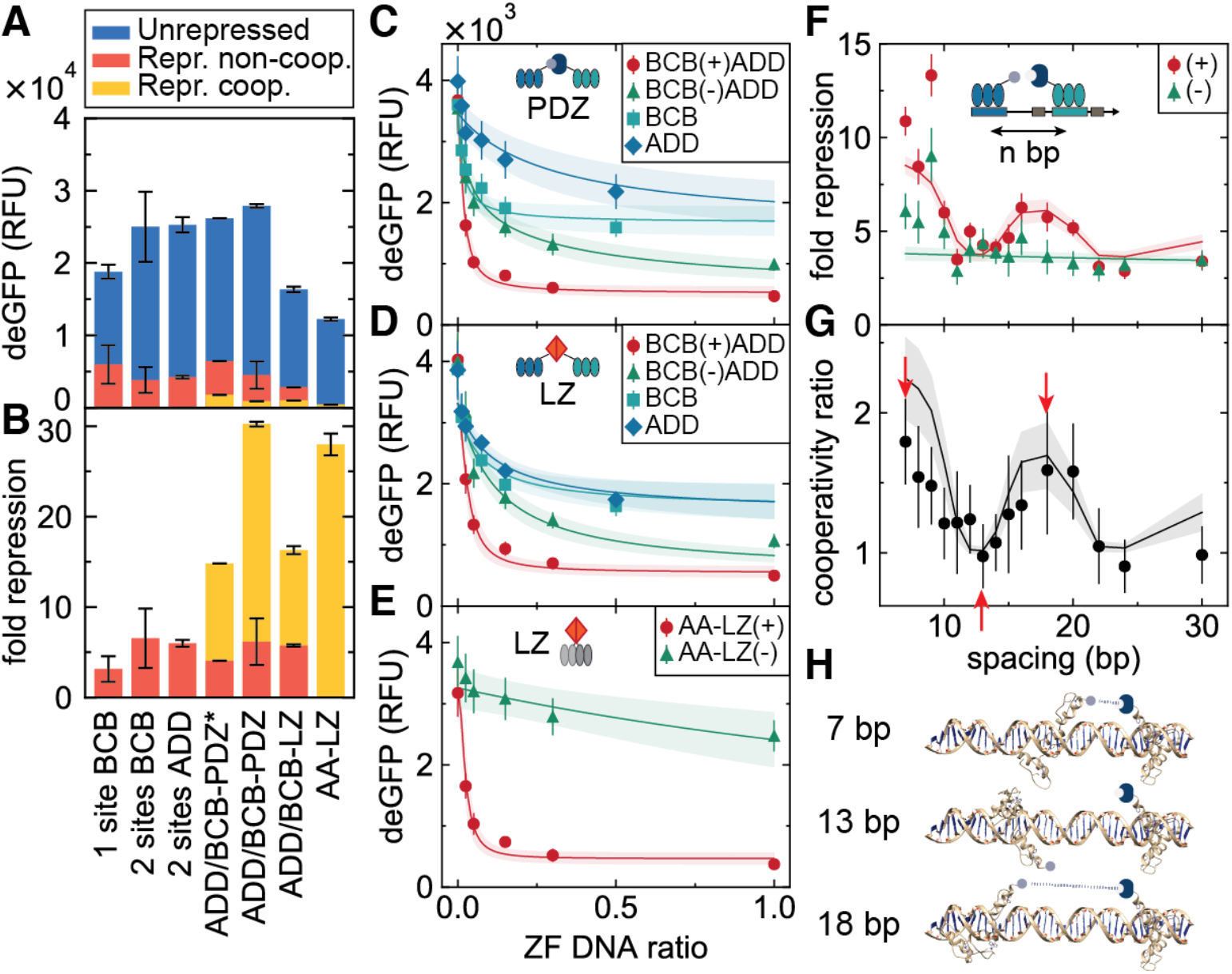
Engineering cooperativity. Comparison of unrepressed and repressed levels (**A**), as well as fold repression (**B**), for different cooperative zinc-finger designs. Three-finger ZFs dimerized using either PDZ-ligand or GCN4 LZ domains. ZFs were linked to interaction domains using proline (-PDZ*) or optimized glycine-serine linkers (-PDZ, -LZ). Additionally, two-finger ZFs were dimerized using LZs (AA-LZ). Data in panels A and B were taken from plate reader experiments; all values represent means ± SD (*n* = 3). On-chip dose response curves for the three-finger PDZ (**C**), LZ (**D**), and two-finger LZ (**E**) designs. The maximum *a posteriori* values as well as 2 SD boundaries of the model predictions are shown as solid lines and shaded regions, respectively. Data in panels C–E was measured on a single chip; all values represent means ± SD (*n* = 12). Shifting the ADD binding site upstream of the −35 box resulted in periodic modulation of the fold repression for the cooperative designs (**F**) as well as in the ratio between the cooperative and non-cooperative fold repressions (**G**), likely due to periodic changes in relative ZF positioning (**H**). All data was collected from a single chip and all values represent means ± SD (*n* = 9).

To investigate cooperativity in more detail, we measured dose response curves by titrating repressor DNA concentration. To keep the load on the transcription-translation machinery constant, the total ZF DNA concentration was kept constant by adding DNA coding for a non-binding ZF control (*ZF_CBD_*). Figure 5C shows dose response curves of *ZF_BCB_ – PDZ* and *ZF_ADD_ – L* separately, together with those for the cooperative pair: *ZF_BCB_ – PDZ + ZF_ADD_ – L*, and the non-cooperative pair: *ZF_BCB_ – PDZ + ZF_ADD_ – NL*. An increase in the steepness of the dose response curve was observed as we proceeded from a single ZF to two non-cooperatively interacting ZFs, and finally to two cooperatively interacting ZFs. Similar results were obtained for the LZ designs (Figure 5D, E). The effect of cooperativity can be quantified by determining the sensitivity (Figure S7), which measures the steepness of the dose response curve (Bintu et al. 2005b), as well as the effective Hill coefficient, which is obtained by fitting phenomenological Hill functions (Figure S8). The results of this analysis are shown in Table S1. We observe that cooperativity increased sensitivity by nearly 50% with respect to the non-cooperative repression, as well as slightly increasing the Hill coefficient.

We sought to understand this behaviour quantitatively by developing a thermodynamic model that relates protein expression to the equilibrium occupancy of the promoter by RNAP (Bintu et al. 2005). We extended the standard competitive model of repression to include a term for the interaction between repressor and RNAP, which is quantified by an effective interaction energy. As this energy tends to large positive values, DNA binding by either RNAP or the repressor is exclusive, and the model tends towards that of competitive inhibition. As the energy approaches zero, both RNAP and DNA can bind simultaneously, resulting in leaky expression at full repressor occupancy. This extension to the model was motivated by our results that a ZF with a fixed binding affinity represses with varying efficiency depending on the position of the binding site; the changing RNAP-ZF interaction energy therefore provides a simple description of this effect. We fit the model to the dose response curves using Markov chain Monte Carlo (MCMC) sampling (Figure S9), allowing us to consistently extract the posterior probability distributions of all parameters, which consist of fixed effective dissociation constants of each individual ZF, as well as the effective energies describing ZF-RNAP and ZF-ZF interactions. The fits are shown in Figure 5C–E as solid lines and shading, which represent the mean and 2 SD boundaries for model predictions, respectively. The values of all fitted parameters are given in Table S2, and a full description of the model is given in the Methods section. We find physically sensible values for all our parameters; in particular, the cooperative interaction energies for PDZ-L (—2.1 ± 0.2 kcal/mol) and LZ (–1.8 ± 0.2 kcal/mol) are consistent with literature values for similar domains (∼ –2 to —10 kcal/mol (Jana et al. 2000, Saro et al. 2007)).

Since the location of the ZF binding site, and hence the relative positioning of ZF and RNAP, is an important determinant of repression efficiency, it is likely that the relative positioning of the *ZF_BCB_ – PDZ* and *ZF_ADD_ – L* binding sites would also determine their ability to interact and subsequently alter their repressive strength. Keeping the *ZF_BCB_ – PDZ* binding site position fixed, we shifted the *ZF_ADD_ – L* binding site further and further upstream. If the two ZFs are positioned on the promoter such that the cooperative PDZ-ligand interaction is unfavorable, we would expect fold repression to be similar to that of the non-cooperative ZFs. In other words, the ratio between the cooperative and the noncooperative fold repression, a quantity we call the cooperativity ratio, should go to unity when the PDZ-ligand interaction cannot occur.

We observed an effect due to this variation of spacing between the two binding sites (Figure 5F), and this behavior corresponded to the relative orientation of the PDZ-ligand domains. As the binding site is shifted, *ZF_ADD_ – L* rotates around the DNA, modulating its alignment with *ZF_BCB_ – PDZ*. The cooperativity ratio fell to 1 when the interaction was unfavorably aligned, but increased again as the domains began to realign (Figure 5G). The cartoon in Figure 5H shows the predicted orientations of the two ZFs as the left-hand site is shifted. The ability of the ZFs to interact over distances of a few tens of bp is likely due to extension of the long flexible glycine-serine linker used to join the *ZF_BCB_* and the PDZ domain. It is unlikely that DNA bending plays a significant role at these distances, due to dsDNA’s much longer persistence length of ∼150 bp.

We incorporated into our model a phenomenological exponential decay of interaction energies with distance, both between the two ZFs as well as between the ZF and the RNAP. Additionally, the ZF-ZF interaction energy was modulated by a periodic function at the frequency of the DNA helical pitch (10.5 bp/turn). Using previously inferred parameters for energies and *K_D_*s from the dose response measurements, we performed a fit to determine the decay constant and phase shift; the results are shown as solid lines and shading in Figure 5F and G, and in Table S2. Fitting a model with an explicit position dependence for the binding sites illustrates the importance of site positioning for functional repression. More generally, while simplistic, our model fits demonstrate that it is possible to understand cell-free gene expression in terms of thermodynamic occupancy.

### E. Affinity tuning

In order to test whether fold repression levels could be precisely and predictively tuned, we investigated the effect of varying binding site affinity. In order to rationally tune binding site affinity, we first generated quantitative PWMs for three ZFs: *ZF_BCB_, ZF_AAA_* and *ZF_ADD_*, covering the nine bp core sequence plus three flanking bases on either side (Figure 6A, S10A, B). The sequence logo determined for *ZF_AAA_* is in concordance with the consensus sequence determined by bacterial one-hybrid and in *vitro* SELEX assays (Meng et al. 2005, Wolfe et al. 1999). Based on our PWMs we designed a library of promoters that included a single binding site at a fixed position between the −35 and −10 boxes, with single or double mutations within or outside the core binding sequence. As binding site affinity decreased we observed corresponding decreases in fold repression for all ZFs tested (Figure 6B, S10C). By converting our macroscopically measured Δ*G* values into microscopic interaction energies Δ*є* we found that the fold repression data could be described by the thermodynamic model presented in the previous section.

**Figure 6:**
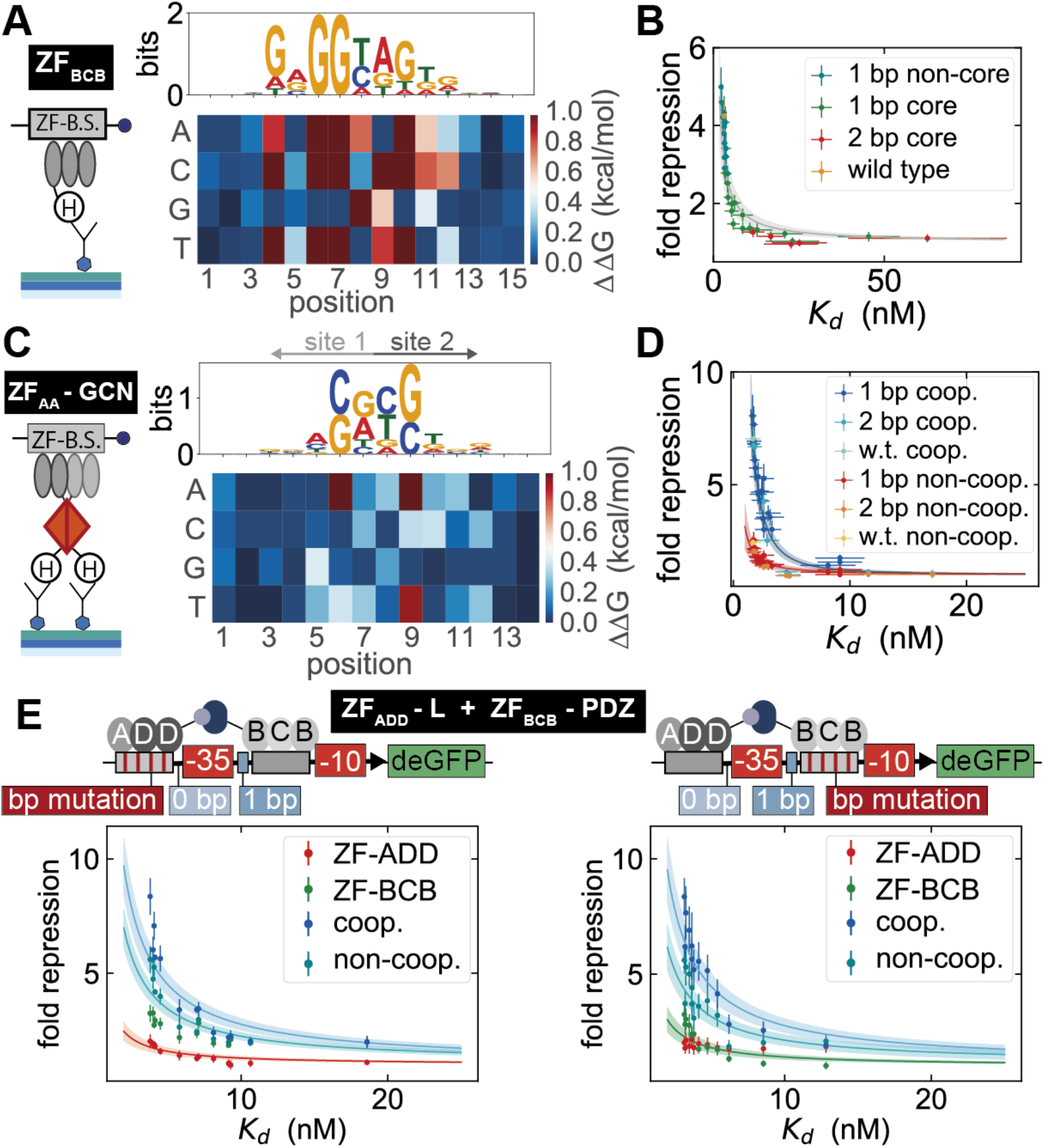
Tuning repression by changing binding site affinity. (**A**) Schematic of the MITOMI assay used for measuring the binding affinity of a ZF to a given DNA target. To the left, the sequence logo and PWM for *ZF_BCB_*, where the core sequence is designated by positions 4-12. (**B**) The relationship between fold repression and *K_d_* values for *ZF_BCB_*. The fold repression data was collected from three separate chips and all values represent means ± SD (*n* = 7). (**C**) Schematic of the MITOMI surface used for measuring the binding affinity of the ZFaa – GCN homodimer to a given DNA target. To the left, the sequence logo and PWM for *ZF_AA_ – GCN*, where the core sequence is designated by positions 3-12. (**D**) The relationship between fold repression and *K_d_* values for both the cooperative and non-cooperative variants of *ZF_AA_ – GCN*. The fold repression data was collected from a single chip and all values represent means ± SD (*n* = 8). (**E**) Fold repression versus *K_d_* values for the *ZF_ADD_ – L - ZF_BCB_ – PDZ* heterodimer pair. On the left the *K_d_*s refer to the *K_d_* arising from the specific change made to the ADD binding site, whereas on the right the *K_d_*s are associated with the BCB binding site. The fold repression data was collected from a single chip and all values represent means ± SD (*n* = 4). In all cases the error bars shown for the *K_d_* values represent the 95 % confidence interval for the fit to a single binding site model, the solid lines are maximum *a posteriori* values from thermodynamic model fits, and the shaded region represents a 2 SD boundary.

Mutating either a single base outside the core site, or one core position of low information content (high entropy), enabled fine tuning of fold repression, whereas a single mutation in the core site of high information content strongly decreased fold repression. Two core mutations decreased fold repression to baseline levels. Fold repression was therefore precisely tuneable over the entire dynamic range by modulating binding site affinity, and the affinity changes required to achieve tuning were relatively small. Affinity changes of ∼ 0.5 to 1 kcal/mol were sufficient to cover the entire dynamic range for each ZF repressor tested. The results are in line with previous findings that promoter tuning in *S. cerevisiae* can be accomplished by relatively subtle affinity changes in a single binding site created by mutations in flanking or single core site mutations of high entropy (Rajkumar et al. 2013). They also correspond to recent results obtained in *E. coli* (Barnes et al. 2018).

Given that a single ZF binding site could be mutated to yield varying levels of repression we investigated whether the same tuning could be applied to cooperative *ZF*s. We measured the binding affinity of the *ZF_AA_ – GCN* homodimer versus a library of DNA targets that consisted of all single point mutations for the 10 bp core binding sequence plus 2 flanking bases on either side. The resulting sequence logo and PWM reveal the symmetric binding profile of the homodimer (Figure 6C). Mutating a single binding site within the −35 and −10 boxes led to a change in repression levels that reflected the measured *K_d_*s for both the cooperative and non-cooperative *ZF_AA_ – GCN* variants (Figure 6D). As the two 6 bp binding sequences overlap, mutating a single base within the core site leads to a finer tuning of fold repression in comparison with the three-finger ZFs. Furthermore, we extended binding site tuning to the *ZF_ADD_ – L - ZF_BCB_ – PDZ* heterodimer pair, taking advantage of the PWMs generated for *ZF_BCB_* and *ZF_ADD_*. Implementing a subset of mutations to each ZF binding site yielded a range of fold repression values not only for the single ZF but also for the cooperative and non-cooperative ZF pairs (Figure 6E). As the affinity of one ZF is reduced we see that the fold repression observed for the cooperative and non-cooperative cases tends to the fold repression measured for the second ZF whose binding site remains constant.

### F. Logic gate construction

Having established a well-characterized resource of transcriptional repressors and promoters, we applied them to designing logic gates. By combining two cooperative ZF repressors on a single promoter we were able to create NAND gates, which are of particular interest as they are functionally complete. An effective NAND gate should have low output only when both inputs are present (Figure 7A). We therefore placed the binding site for a strongly binding ZF (*ZF_BCB_*) 2 bp upstream of the −35 box, and second binding site for different ZFs between the −35 and −10 boxes. *ZF_BCB_* cannot strongly repress by itself at the −2 position and the second ZF should also not strongly repress on its own. Only when both ZFs are bound to the promoter should they strongly repress, which can be achieved by including a cooperative interaction between the two ZFs. Using this general design we tested NAND gates for *ZF_BCB_ – PDZ* in combination with the remaining ZFs (Figure 7B). As expected, NAND gate performance improved as the affinity of the *ZF_XXX_ – L* decreased. For instance the combination of *ZF_BCB_ – PDZ* and *ZF_BDD_ – L* gave rise to a functional NAND gate, whereas a combination with *ZF_AAA_ – L* did not due to the high affinity of *ZF_AAA_ – L*, which led to functional repression even when only *ZF_AAA_ – L* was present.

**Figure 7:**
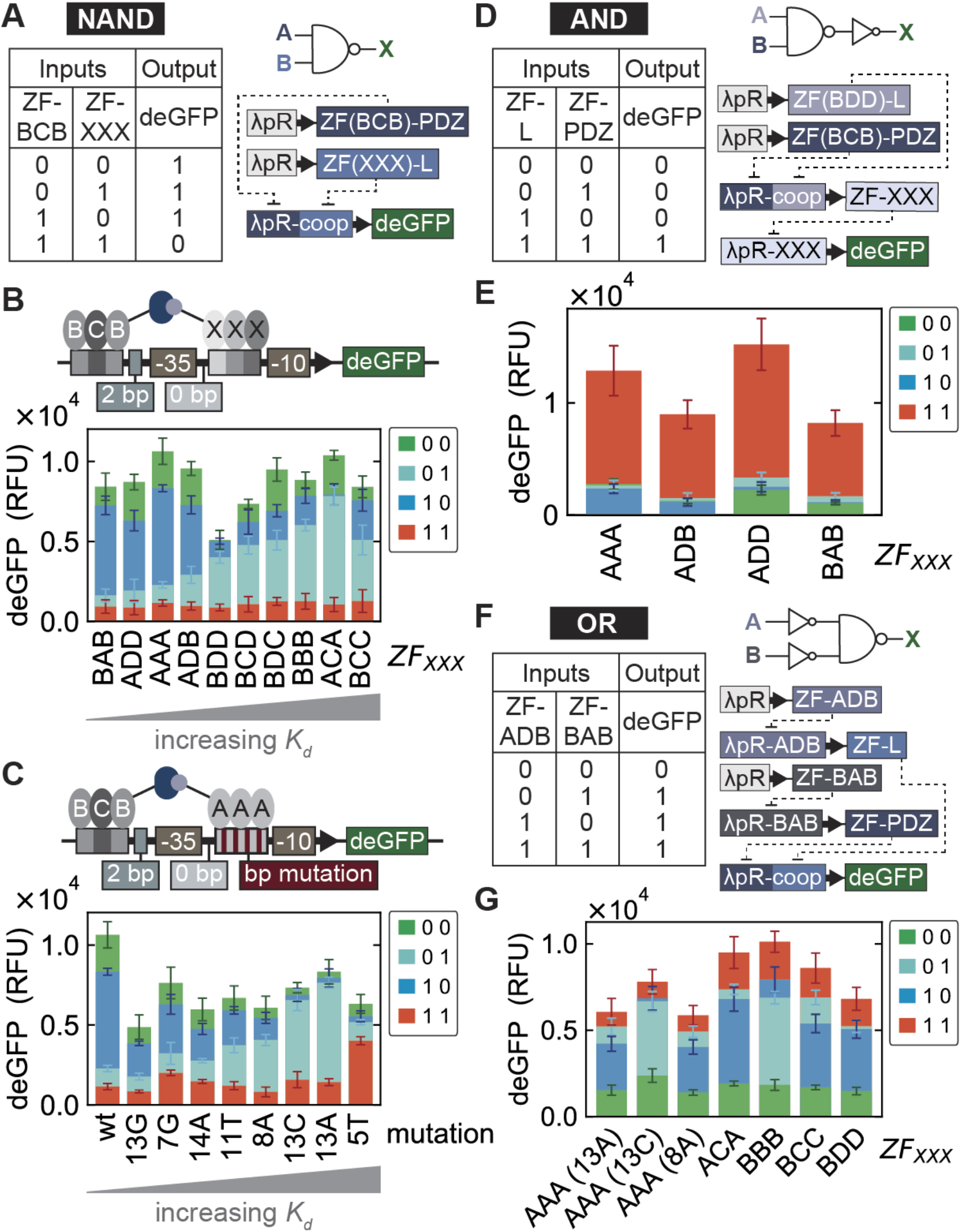
Logic gates. (**A**) Truth table, logic gate symbol and biological network design for constructing a NAND logic gate. (**B**) NAND gate design combining *ZF_BCB_ – PDZ* with all other *ZF_XXX_ – L*s. Below, the output for each NAND gate tested. (**C**) The same NAND gate design as in (**A**) except that only *ZF_BCB_ – PDZ* and *ZF_AAA_ – L* are used as inputs and the *ZF_AAA_ – L* binding site affinity is rationally adjusted to yield a functional NAND gate. The deGFP outputs for all NAND gates were measured from a single chip where all values presented correspond to the mean ± SD (*n* = 5). (**D**) Truth table, logic gate symbols and biological network design for constructing an AND logic gate. (**E**) The output for each AND gate tested. All output values were measured from a single chip and represent the mean ± SD (*n* = 8). (**F**) Truth table, logic gate symbols and biological network design for constructing an OR logic gate. (**G**) The output for each OR gate tested. When *ZF_AAA_ – L* is used as part of the NAND gate, the mutation of the binding site is indicated in the parentheses. All output values were measured from a single chip and represent the mean ± SD (*n* = 6).

Since we showed that binding affinity could be precisely tuned (Figure 6) we tested whether we could improve our non-functional NAND gates. Based on the PWM measured for *ZF_AAA_* we mutated the *ZF_AAA_ – L* binding site sequence in the NAND gate promoter and showed that we could achieve tuning in this context as well (Figure S10D). We then investigated the effect of tuning the *ZF_AAA_ – L* binding site for all possible input combinations and showed that the NAND gate improved as we weakened *ZF_AAA_ – L* binding affinity (Figure 7C). Mutations +1C and +1A gave rise to functional NAND gates. Decreasing the binding site affinity increased the output when only *ZF_AAA_ – L* was present; however, when the mutation resulted in a ΔΔ*G* of greater than ∼ 0.5 kcal/mol (Δ2*T*), the cooperative binding output also suffered. Our synthetic ZF repressors can thus be used to build functional NAND gates, which can additionally be rationally optimized and precisely tuned by modifying binding site affinities.

As a final example we generated compound logic gates by combining NAND and NOT logic gates as linear cascades in order to create AND and OR gates. We created an AND gate by appending a NOT gate to the output of a NAND gate (Figure 7D). Specifically we combined the *ZF_BDD_ – L - ZF_BCB_ – PDZ* NAND gate with four different ZFs. Each AND gate was tested and yielded the expected outputs (Figure 7E). We then generated OR logic gates by prepending two NOT gates in front of different NAND gates to invert the inputs (Figure 7F). We used *ZF_ADB_* and *ZF_BAB_* as the two NOT gate inverters and a set of NAND gates, all of which gave rise to functional OR gates (Figure 7G).

## DISCUSSION

GRNs are of central importance in both native and engineered systems. They integrate, compute, and transduce input signals, leading to specific changes in gene expression. Many components contribute to the function of GRNs, and transcription factors and their interaction with promoters are core players. Due to the complexity of even a single transcription factor – promoter interaction it has proven difficult to quantitatively study these systems *in vitro* or *in vivo*. Although the development of new technologies is steadily enabling progress in this area, our understanding of GRNs remains limited as exemplified by our inability to predict *in vivo* gene expression levels in essentially any organism, and the difficulty asso ciated with *de novo* engineering of GRNs. Although methods exist for high-throughput in *vitro* characterization of transcription factor binding specificities (Bulyk et al. 2001, Jung et al. 2018, Maerkl and Quake 2007, Zhao et al. 2009) and medium to high-throughput approaches are used to understand gene regulation *in vivo* (Barnes et al. 2018, Mogno et al. 2013, Rajkumar et al. 2013, Sharon et al. 2012) both approaches have limitations. Both an advantage and disadvantage of *in vitro* methods is that they generally include only the smallest number of components necessary, i.e. a transcription factor, dsDNA target and a defined buffer solution. *In vivo* methods are on the other hand convoluted by cellular complexity. Furthermore, generating and analyzing defined libraries *in vivo* remains labor intensive and difficult. Here we explored the use of a cell-free transcription-translation system to build and characterize GRNs in an environment that bridges the gap between *in vitro* and *in vivo* methods. This cell-free approach also has the advantage of allowing complex assays to be performed in high-throughput, in a well-controlled and accessible environment. As a consequence, the ability to study functional transcriptional regulation in an *in vitro* system has allowed us to delve into much greater depth than comparable *in vivo* methods have been able to achieve (Amit et al. 2011, Garcia and Phillips 2011, Rajkumar et al. 2013)

We chose to build GRNs from the bottom up using ZF transcription factors for several reasons. First, in regards to GRN engineering, researchers have long been hampered by the relatively small number and poor characterization of available transcriptional regulators. Khalil *et al*. have previously engineered ZF regulators, showing that they are viable tunable transcriptional regulators *in vivo* (Khalil et al. 2012). We built on this concept, generating additional ZF regulators and interaction domains. More importantly, we quantified the binding energy landscapes of several synthetic ZF regulators and were able to show that repression can be precisely tuned with small changes in affinity. These small changes were achieved by mutating the flanking bases lying outside of the consensus core sequence or by mutating one consensus core base of low information content. Hitherto, only coarse tuning has been accomplished through varying the number of consensus sequence binding sites leading to rather large differences in output (Khalil et al. 2012, Lohmueller et al. 2012). The ability to predictively and precisely tune expression levels as demonstrated here is important in engineered GRNs where individual nodes of the network need to be matched in expression levels. For example, we show here that the ability to precisely adjust individual binding site affinities is crucially important for optimizing logic gate function.

With the advent of TALEs and dCas9, ZFs might be considered outdated technology, but there are a number of reasons why ZF TFs remain an appealing tool for GRN engineering. ZFs have several advantages such as small size, relatively easy gene synthesis, and good expressability. The biggest advantage of dCas9 and TALEs is their programmability, allowing them to be precisely targeted to any DNA sequence. Conversely for ZFs, it remains relatively difficult to rationally design a particular binding site preference. For genome editing and *in vivo* targeting approaches, in which the target sequence is defined and immutable, programmability is crucial. In the context of bottom up GRN design, this ability becomes less important as target sequences can be easily adjusted to a particular TF specificity. We argue that it is actually more important to be in possession of a well-characterized TF binding energy landscape that can be obtained for ZF TFs using current methods (Blackburn et al. 2015).

A second argument in support of using ZF transcription factors over TALEs and dCas9 is the simple but important fact that ZFs are native transcriptional regulators and the most abundant class of transcriptional regulators *in vivo*. Cas9, to the best of our knowledge, has not been shown to be involved in gene regulation in native systems, while TALEs are injected into plant host cells to modulate gene expression by pathogenic bacteria (Boch et al. 2009). If cell-free approaches are to be used to understand the function of native systems it is important to build GRNs with native transcription factors. For example, the protein – DNA interaction kinetics are very different in that dCas9 (Boyle et al. 2017) and TALE (Cuculis et al. 2016) tend to have very slow DNA dissociation rates, while native transcriptional regulators have fast dissociation rates (Geertz et al. 2012), which may make engineering dynamic GRNs using TALEs and dCas9 difficult.

In order to improve fold repression and to add more control over the system we engineered cooperative binding into our ZF TFs by including PDZ or LZ protein – protein interaction domains. These interactions improved repression from ∼10 to up to ∼30 fold and were functional for both two- and three-finger ZFs. We showed that the relative placement of binding sites for two cooperative TFs is a major determinant of interaction capacity and consequently repression strength. Repression was achieved when the TFs were located on the same face of the DNA, and repression strength followed the helical twist of DNA. Cooperative interactions consequently allowed us to engineer functionally complete NAND gates. In all cases we were able to explain our data with thermodynamic models. Combining these models with binding energy landscapes thus provides a viable and useful approach to rationally engineer GRNs.

One outstanding problem encountered during this study is the issue of composability. Although transcription factor binding sites were only introduced in regions outside the −10 and −35 boxes of the original *λP_R_* promoter, many of the synthetic promoters had considerably different baseline (non-repressed) expression levels. In the future it will clearly be important to better understand and predict basal promoter strength from the underlying sequence, which would lead to models that allow introduction of transcription factor binding sites without affecting basal promoter output. Here we have seen that basal promoter strength itself can be finely tuned over a relatively large range of expression levels (Figure 1). It should therefore be possible to adjust promoter strength as desired: we demonstrate a basic example of this idea by tuning the basal expression level of a repressible promoter (Figure S11). Ultimately understanding the outcome of multiple base changes in close context with each other remains a complex issue. Evaluating a greater number of sequences and systematically addressing all factors affecting transcription efficiency similar to the approach taken by Cambray *et al*. towards translation could lead to an improved understanding of promoter sequence design principles (Cambray et al. 2018).

In order to characterize and measure our synthetic ZF transcription factors and promoters in detail we repurposed a high-throughput microfluidic device that allowed us to measure 768 cell-free reactions in parallel. Eliminating cloning and transformation steps by relying on PCR-based assembly strategies allowed us to measure a large number of defined transcription factor and promoter variants. Over 13,000 on-chip cell-free TX-TL reactions were performed, encompassing replicates for ∼2000 unique reactions. We furthermore took over 8000 MITOMI measurements to provide binding energy landscapes for 4 synthetic ZF transcription factors. Together, these technologies allowed us to establish a quantitative and in-depth dataset and insights into transcriptional regulation that should be of general interest. The approach taken here nonetheless does not *per se* require these state-of-the-art technologies, and is easily transferable to standard lab equipment. Cell-free lysate can now be easily and cheaply generated, yielding sufficient material so that medium-scale screens in 384-well plates are feasible (Sun et al. 2013). Commercial liquid handling equipment can also be used to scale up throughput. Binding energy landscapes can be generated by many approaches including PBMs (Bulyk et al. 2001), MITOMI (Maerkl and Quake 2009), SELEX-seq (Zhao et al. 2009), and HiP-FA (Jung et al. 2018). While our binding energy landscapes are based on direct affinity measurements, it may be sufficient to use PWMs from indirect measurements as found in other high-throughput techniques.

Rapid progress is being made in the development and application of cell-free synthetic biology. Cell-free systems are being used to tackle fundamental problems in molecular engineering and are being applied to molecular diagnostics (Pardee et al. 2016), therapeutics (Pardee et al. 2016b), synthesis (Goering et al. 2016), and are even being used for educational purposes (Stark et al. 2018). Cell-free systems are an appealing alternative to cellular systems, as they eliminate many of the complexities associated with working with cells. Cell-free systems are also a rapid prototyping platform for engineering molecular systems destined to be applied in cellular hosts (Niederholtmeyer et al. 2015). As engineered systems become more complex it will become increasingly important that a large number of standardized characterized components become available. It will be equally important to develop a comprehensive mechanistic understanding of these components and systems to allow parts to be standardized and rationally assembled without requiring extensive trial-and-error cycles or large screens, which may not be feasible for large systems. As work progresses on cellular sub-systems such as gene regulation, DNA replication, ribosome biogenesis, metabolic networks, and membrane and protein super-structures, it will be intriguing to contemplate whether it may be possible to integrate these individual systems to create a synthetic cell or cell-like mimic. Work in this area will not only provide tools and methods aiding engineering of synthetic systems, but is likely to provide insights into the function of native systems as well. Prior to being used as tools for protein synthesis and synthetic biology, cell-free systems have already had a rich history in deciphering fundamental aspects of biochemistry including DNA replication (Fuller and Kornberg 1983) and the genetic code (Nirenberg and Matthaei 1961). It is likely that they will continue to provide fundamental insights into complex systems such as transcriptional regulation.

## ACKNOWLEDGEMENTS

We thank Samuel Clamons and Miki Yun from the Murray lab (Caltech) for providing the cell-free transcription-translation extract and Samuel Clamons and Richard Murray for helpful discussions. We also thank Malek Kabani, Eugenia Pankevich, and Stefan Bassler for their experimental contributions to this project. This work was supported by an HFSP Program Grant (RGP0032/2015) and the École Polytechnique Fédérale de Lausanne.

## AUTHOR CONTRIBUTIONS

Z.S. and N.L. performed experiments. Z.S., N.L. and S.J.M. designed experiments, analyzed data and wrote the manuscript.

## SUPPLEMENTAL FIGURES

**Figure S1:**
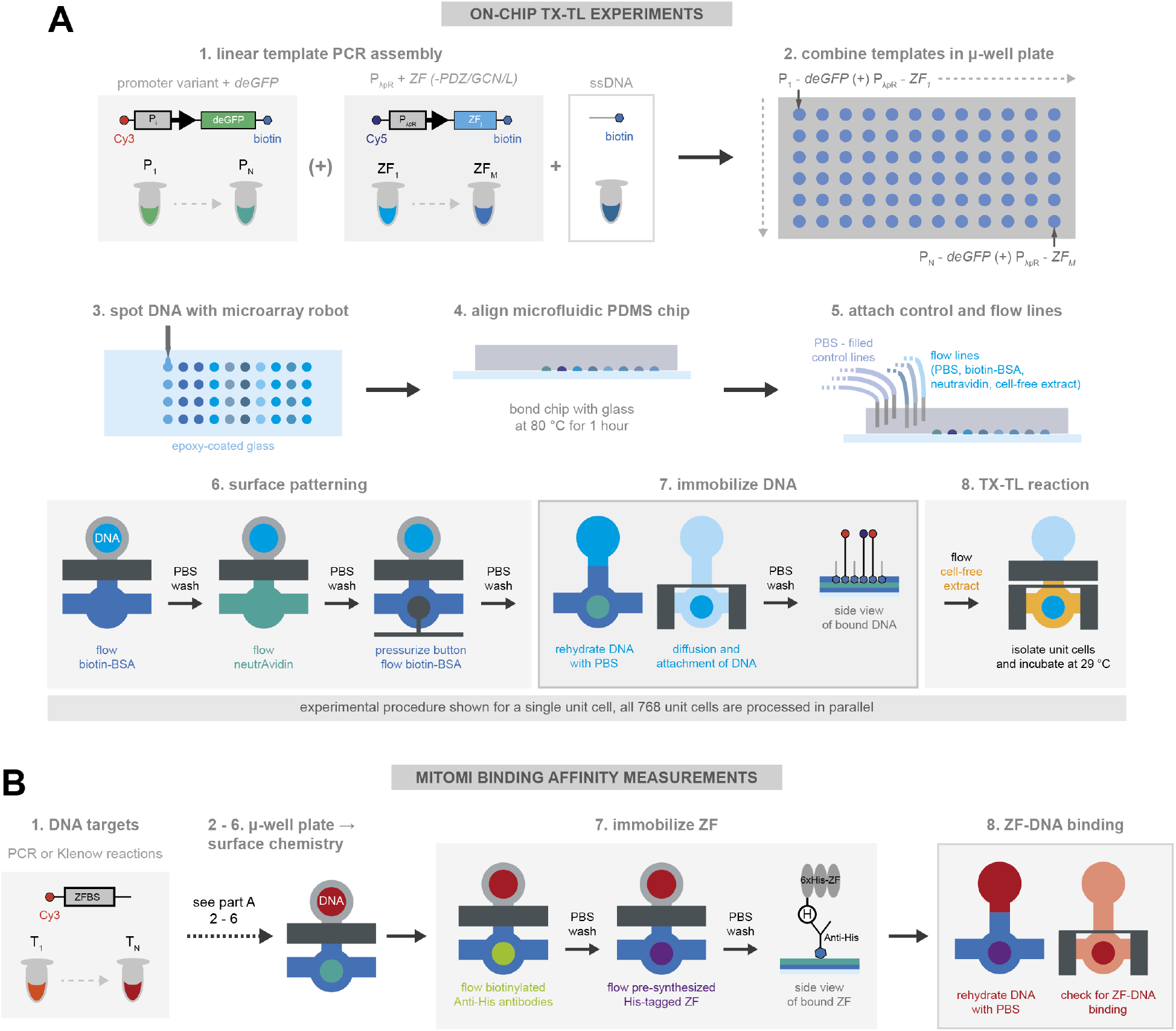
Overview of experimental protocol. Schematic showing the details of the experimental procedure for on-chip cell-free TX-TL reactions (**A**) and for measuring the binding affinity of ZF-DNA complexes with MITOMI (**B**).

**Figure S2:**
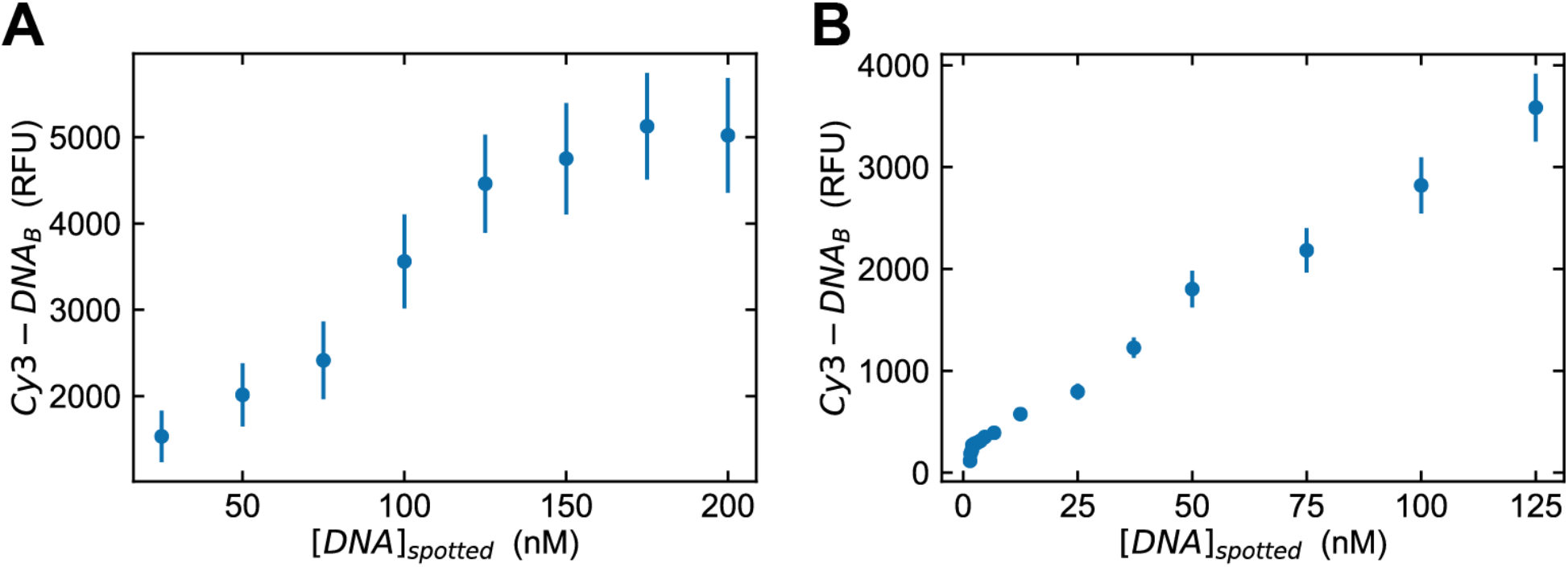
DNA template spotting. DNA concentration measured on-chip versus the concentration of DNA in the spotting plate, when a single dsDNA template was diluted (**A**) versus a mixture of dsDNA template and ssDNA (**B**).

**Figure S3:**
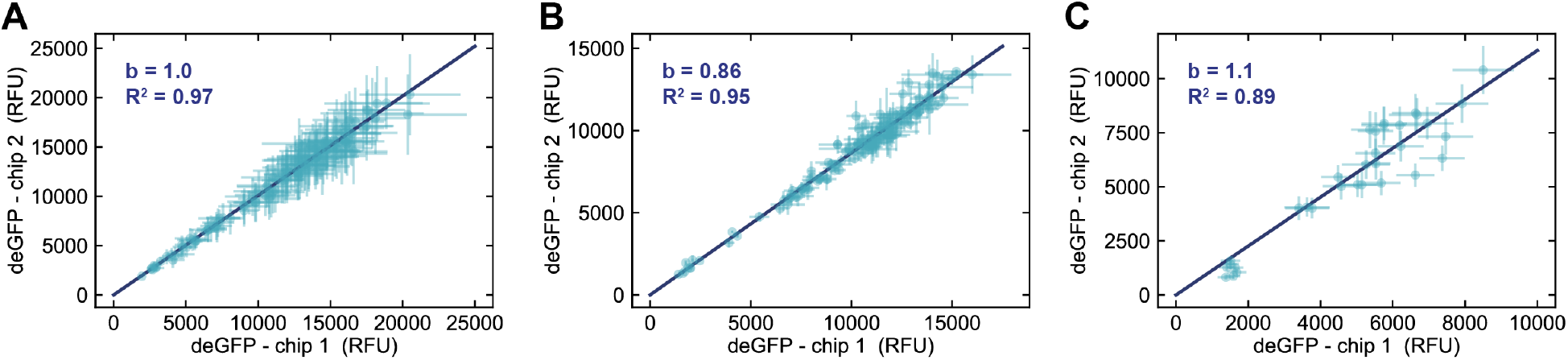
Chip-to-chip reproducibility. (**A**) deGFP expression values obtained from two separate chips for the λ*P_R_* promoter mutagenesis library. The same DNA microplate was used to spot both chips. (**B**) A comparison of two separate chips measuring the output of all possible ZF – promoter pairs. Similar to when only a single DNA template is present per unit cell ((**A**)), two templates could be added with good reproducibility between chips. (**C**) Data from two separate chips with two or more DNA templates per unit cell (NAND logic gate). Each chip was prepared using different DNA microplates, showing a slightly increased variability for DNA templates derived from different PCR reactions.

**Figure S4:**
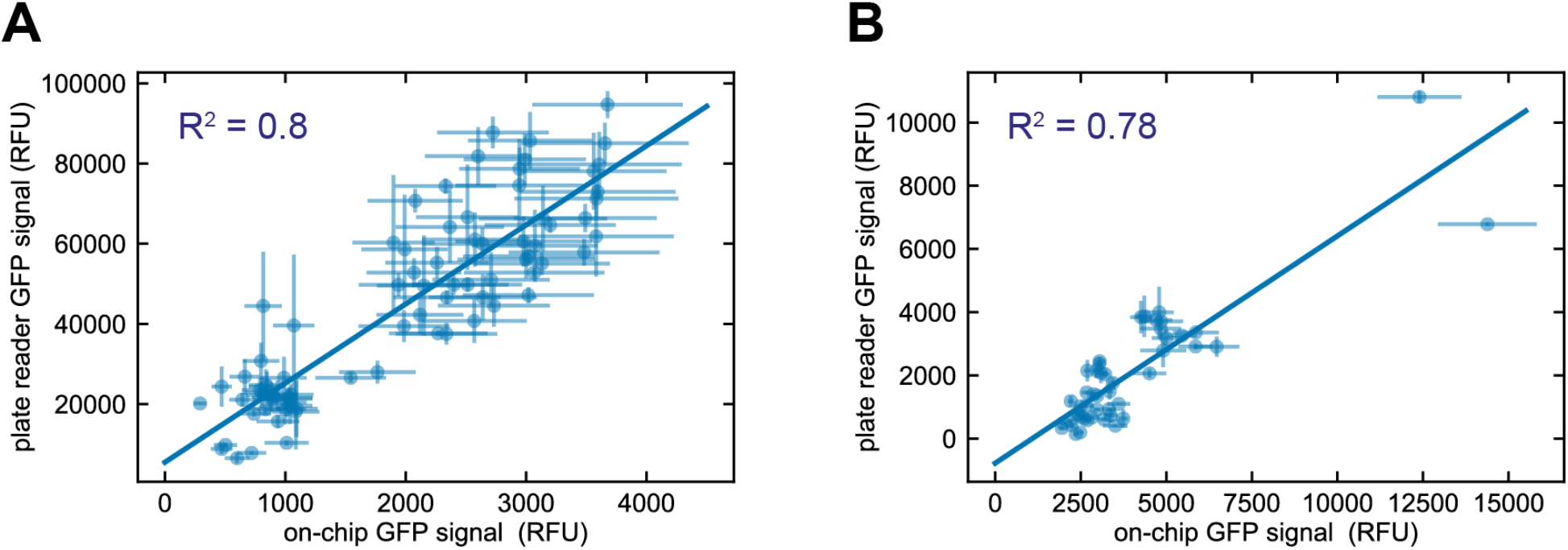
Comparison of on-chip and plate reader measurements. (**A**) Data collected from standard micro-well plate reader reactions versus data collected on-chip for a 9×9 ZF-target orthogonality matrix for targets with two binding sites in the promoter region. The 9 ZFs tested included AAA, ADB, ADD, BAB, BCB, BCC, BCD, BDD and CBD. (**B**) Plate reader versus on-chip data for cooperative, non-cooperative and control ZF PDZ-L heterodimers tested with a library of promoters that had variable spacing between the two ZF binding sites.

**Figure S5:**
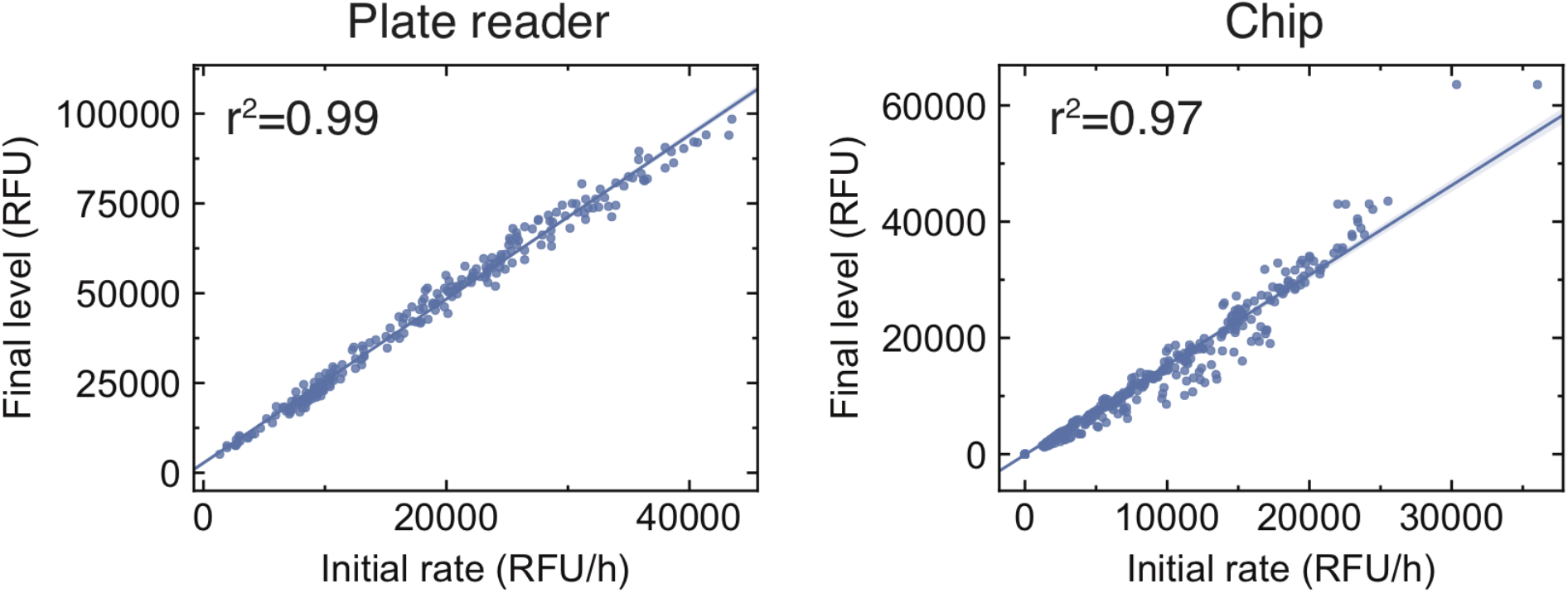
Relationship between initial rates and final deGFP values. We observe a linear relationship between the initial rate of deGFP production, as obtained by linear fits to the time course at early times (∼30 minutes), and the final steady state level of deGFP, on both plate reader and chip experiments. This suggests that final levels of deGFP are proportional to the production rate, and validates the use of endpoint protein levels as a proxy for transcription rates.

**Figure S6:**
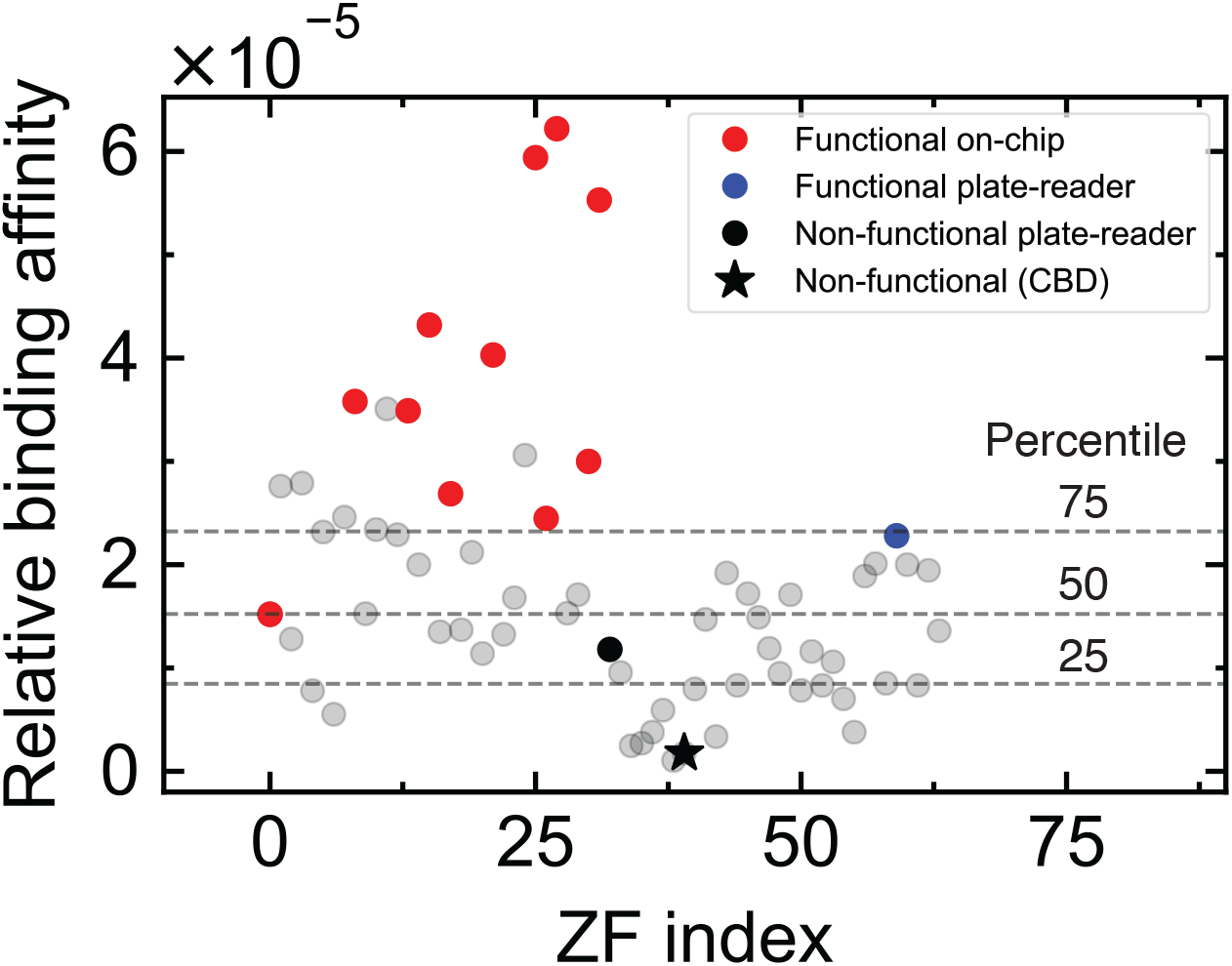
Functional ZFs within the combinatorial library. This figure shows the on-target binding affinity for all 64 members of our combinatorial library (data from (Blackburn et al. 2015)). In red are the 11 functional ZFs characterised on-chip. Additionally, the blue and black points are two other ZFs characterised in a separate plate-reader experiment. The black star corresponds to *ZF_CBD_*. Horizontal dashed lines represent the 25th, 50th, and 75th percentile of binding affinity, and we observe that of the ZFs tested, all those with binding affinities greater than the 50th percentile exhibited functional repression. We therefore hypothesize that of the untested ones, those with affinities greater than approximately 1.5 × 10^−5^ in relative units should be functional. This would correspond to a total of 32 functional repressors.

**Figure S7:**
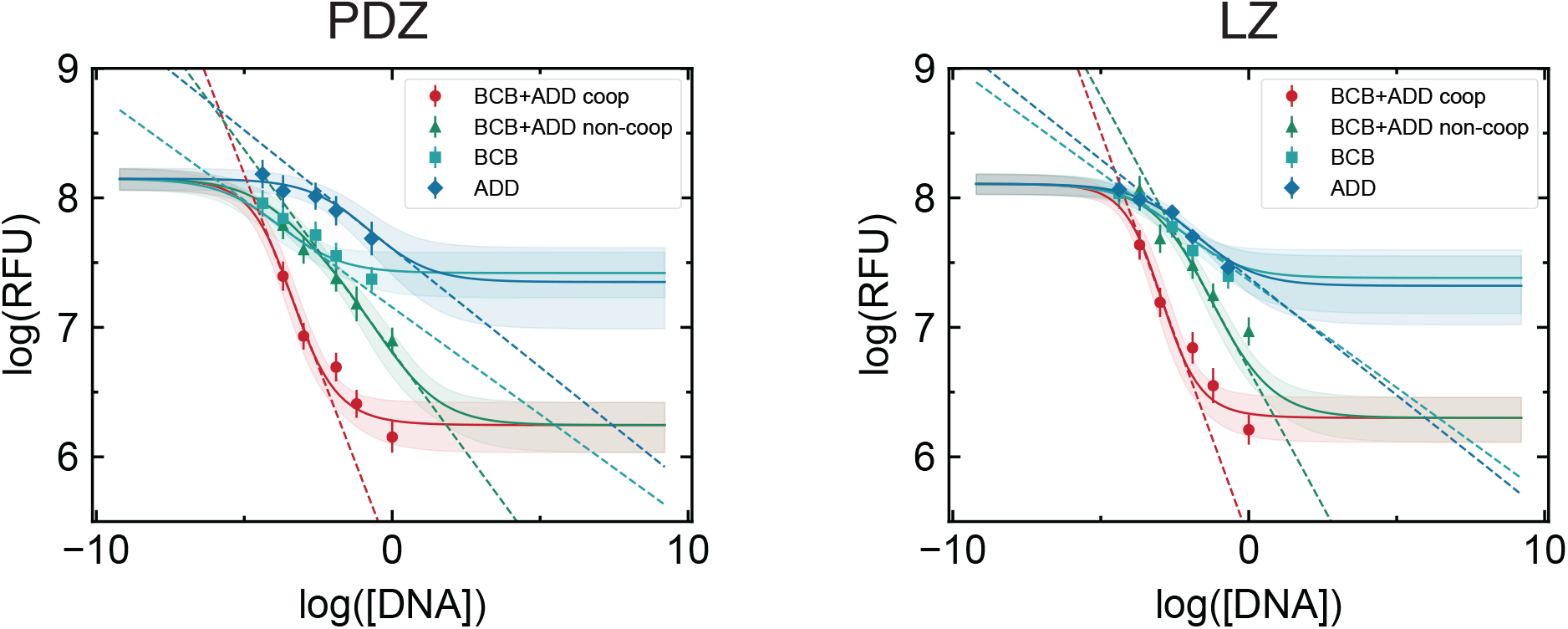
Sensitivity calculations from dose response curves. The slope of the dose response curve, as measured in the linear regime of a log-log plot, is defined as the sensitivity; this quantity increases in the presence of cooperative interactions. Values are given in Table S1.

**Figure S8:**
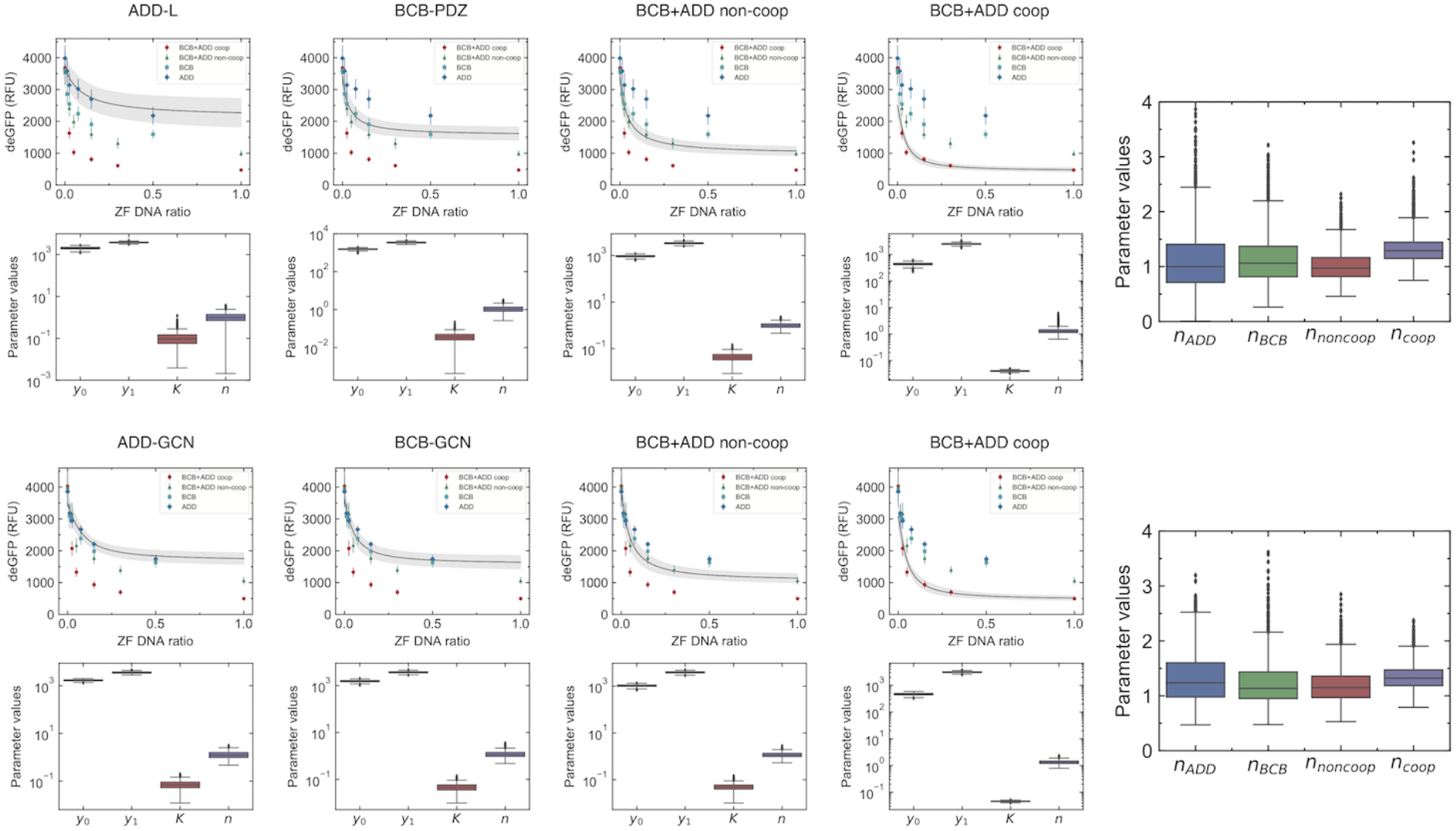
Hill function fits to dose response curves. Standard Hill functions take the form 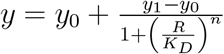, where *y*_0_ is the leak, *y*_1_ the maximum expression, *R* the repressor concentration, *K_D_* the dissociation constant, and *n* the Hill coefficient. Unlike the thermodynamic model which proposes a mechanism to consistently fit the entire data set, Hill functions must be independently fit to each dose response curve. We first fit the single ZF data, followed by the non-cooperative BCB+ADD curve. Hill functions describe two-site binding using an effective *K_D_* and varying *n*. We make the assumption that in both the cooperative and non-cooperative case, the effective *K_D_* is the same. Thus the cooperative BCB+ADD curve is fit using the *K_D_* value obtained from the non-cooperative BCB+ADD curve. Although the Hill coefficient is predicted to increase in the presence of cooperativity, we observe a minimal change in our data, within the error of the fits.

**Figure S9:**
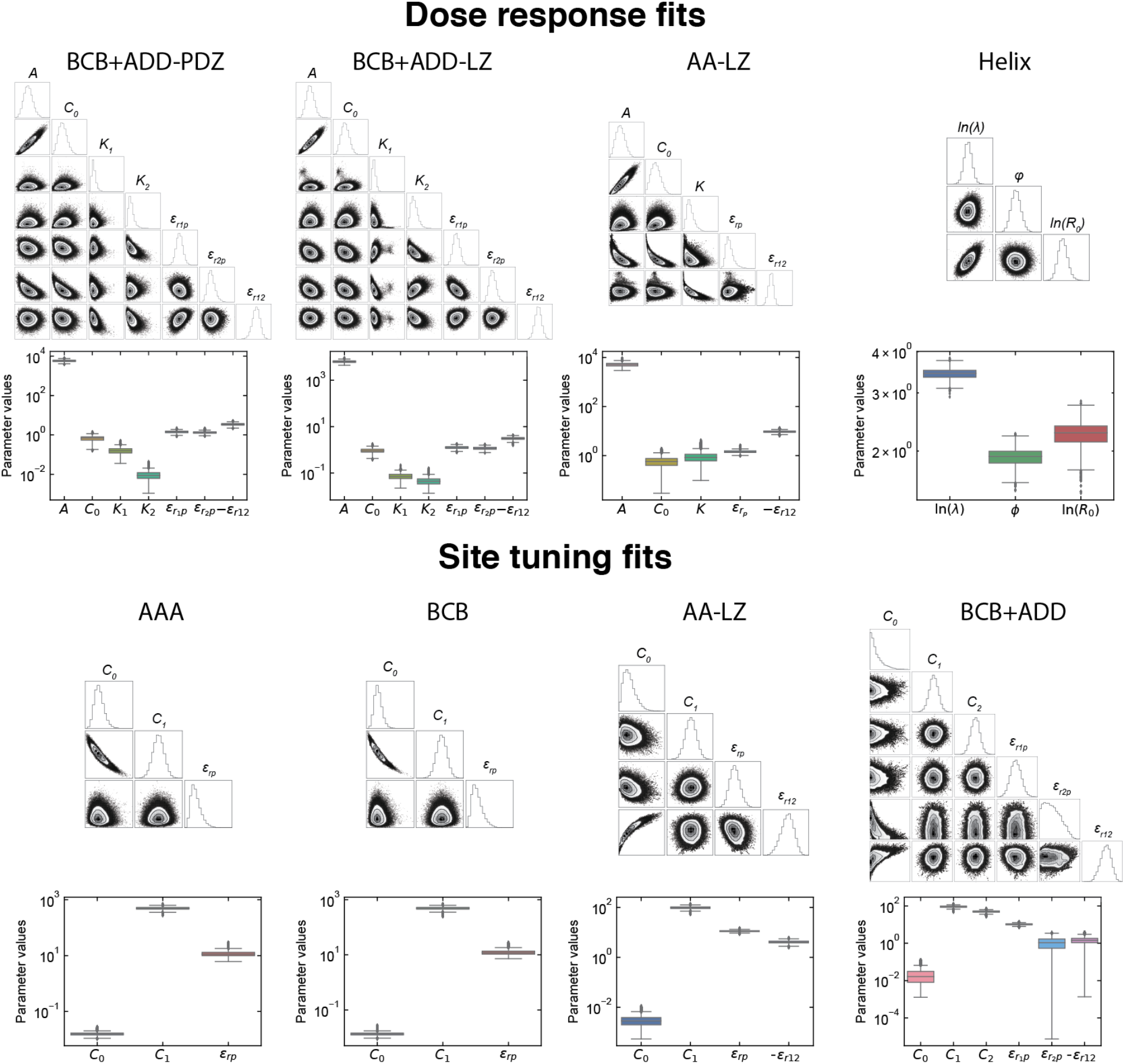
Markov chain Monte Carlo inference of model parameters. Matrix plots show pairwise posterior probability distributions of model parameters for each of the models fitted. The individual posterior distribution for each parameter can also be visualized using box plots; the top and bottom of each box represents the 75th and 25th percentile, respectively.

**Figure S10:**
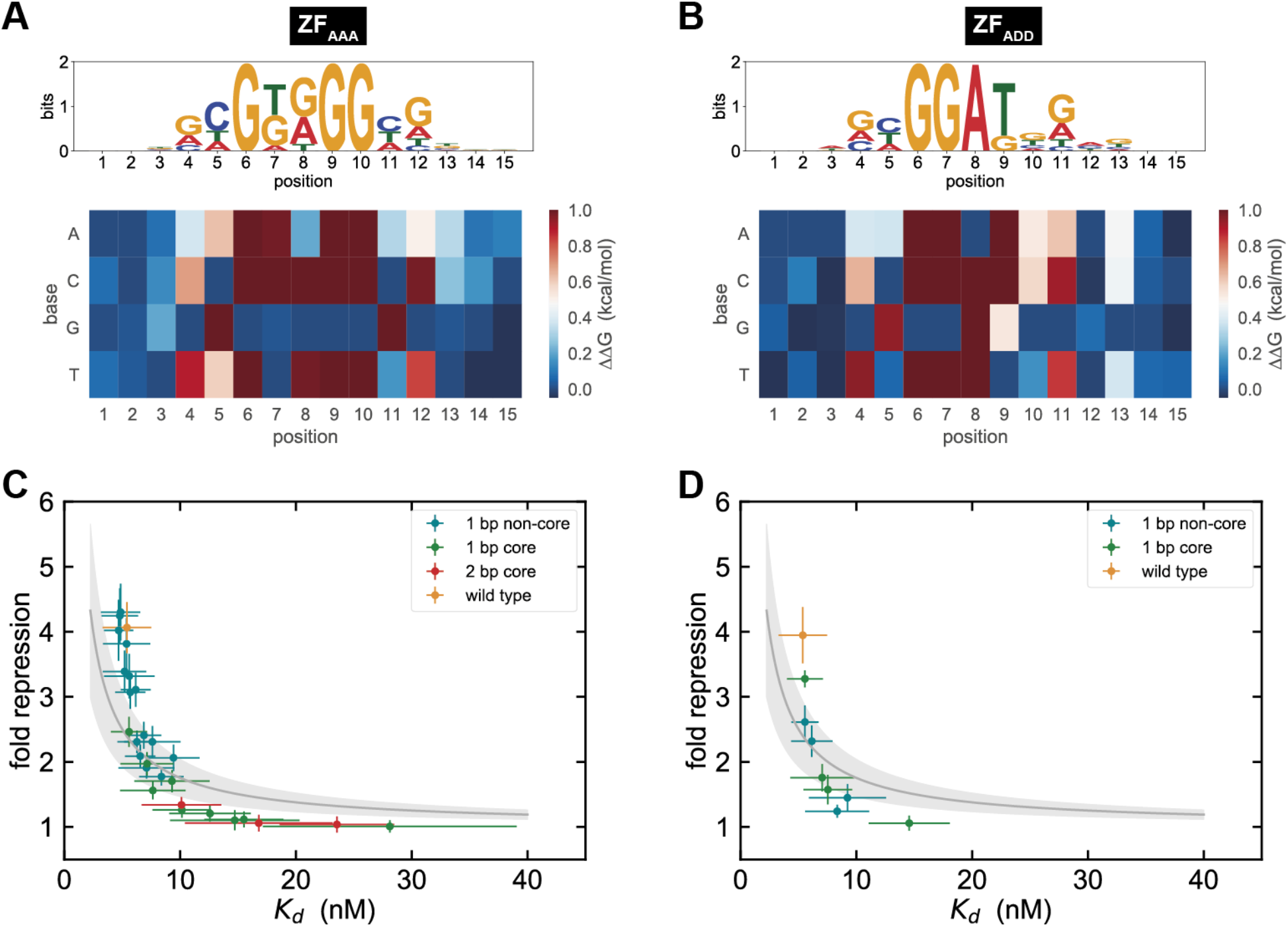
PWMs *ZF_AAA_* and *ZF_ADD_* and binding site tuning for *ZF_AAA_*. Sequence logos and PWMs measured by MITOMI for *ZF_AAA_* (**A**) and *ZF_ADD_* (**B**). Fold repression versus *K*_*d*_ measured for mutations to a *ZF_AAA_* binding site within a single binding site promoter (**C**) and a cooperative promoter (**D**). Solid lines represent the model fits.

**Figure S11:**
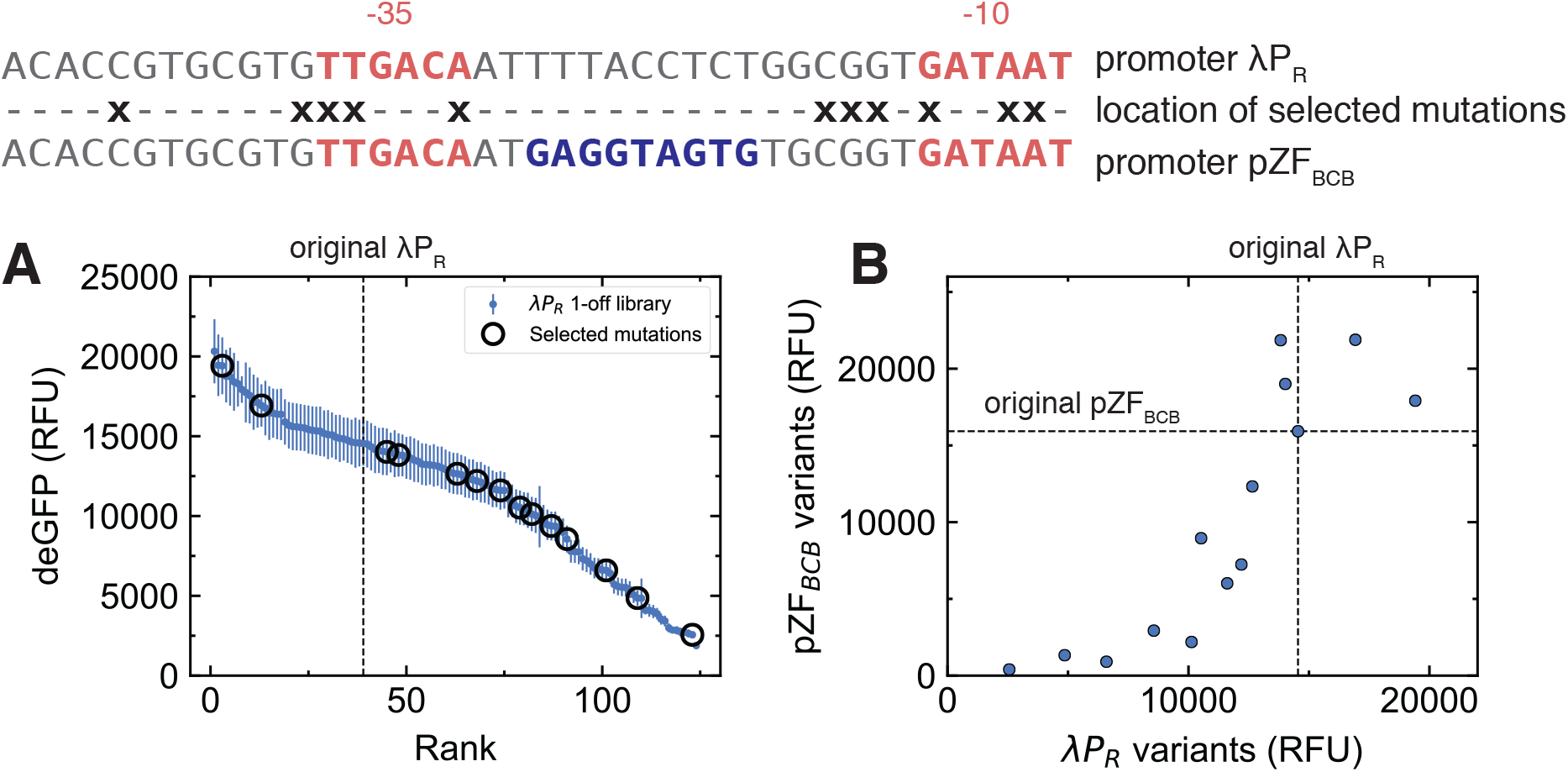
Tuning promoter strength. In Figure 1 we presented the one-off library data which contains all single base mutations from positions −47 to −7 of the λ*P_R_* promoter. (**A**) These data can be rank-ordered, and we observe that a large dynamic range of promoter output is accessible. A number of mutations were selected which both increased and decreased expression relative to the original promoter. (**B**) Introducing a single *BCB* binding site between the −35 and −10 boxes generates the *pZF_BCB_* promoter. Applying the selected mutations to this new promoter changes the output in a correlated way when compared to the λ*P_R_* promoter; the new promoter can thus be roughly tuned in a predictive fashion.

## SUPPLEMENTARY METHODS

### Microfluidic chip fabrication

The molds for each device layer were fabricated using standard photolithography. For the control layer, a silicon wafer was primed in an oxygen plasma processor for 7 minutes (TePla 300) and SU-8 photoresist (GM 1070, Gersteltec Sarl) was spin coated onto the wafer yielding a height of 30 *μ*m. Following a soft bake the wafer was exposed to light (365 nm illumination, 20 mW/cm^2^ light intensity) using a chrome mask for 10 s on a Süss MJB4 mask aligner. After a post exposure bake the wafer was developed with PGMEA (propylene glycol monomethyl ether acetate) to remove unexposed SU-8 and a hard bake was performed to remove unwanted cracks in the SU-8 structures. For the flow layer, a silicon wafer was treated with HMDS (hexamethyldisilazane) vapor in a YesIII primer oven and AZ 9260 photoresist (Microchemicals GmbH) was spin coated on the wafer to a height of 14 μm. After baking and a one hour relaxation period, the coated wafer was illuminated with a broadband light using an MABA6 mask aligner with a total dose of 660 mJ/cm split into two exposures of 18 s with a 10 s wait period between (20 mW/cm^2^ light intensity). The wafer was developed with AZ 400K developer and baked at 175 °C for two hours to re-flow and anneal the AZ structures.

Each of the wafers was subsequently treated with TMCS (trimethylchlorosilane) and coated with PDMS (polydimethylsiloxane, Sylgard 184, Dow Corning). For the control layer PDMS with an elastomer to crosslinker ratio of 5:1 was prepared and poured over the wafer to yield a height of ∼0.5 cm, while for the flow layer, PDMS with a 20:1 elastomer to crosslinker ratio was spin coated at 1800 rpm to yield a height of ∼50 *μ*m. Both PDMS coated wafers were placed in the oven at 80 ^°^C for 20 minutes. After an initial curing the control layers were cut out and the holes for each control line were punched using a 900 mm pin. Each control layer was then aligned by hand on top of a flow layer using a Nikon stereo microscope and the aligned devices were placed in the oven at 80 °C for 90 minutes, allowing the two layers to bond together.

### Preparation of cell-free extract

BL21 Rosetta *E. coli* cell-free extract was a gift from the Murray lab (Caltech) and prepared according to a published protocol (Sun et al. 2013). As described in the citepd protocol, the addition of purified gamS protein is added to the final reaction mixture to prevent the degradation of linear DNA templates by nucleases.

### Preparation of DNA templates

All promoters were generated by PCR from a pBEST-OR2-OR1-Pr-UTR1-deGFP-T500 plasmid (Shin and Noireaux 2010) (Addgene #40019). Oligos containing the promoter variants were ordered from IDT. Assembly PCR was used to synthesize the complete template including a given promoter variant and the downstream deGFP gene. In the first step two linear DNA products are amplified with PCR where each product contains a 20 bp overlap region and one of the products contains the change to the base promoter. The overlap region was generally between the −10 box and the RBS. In the second step the two DNA products were annealed together and the entire template was amplified using global primers which were labeled with a fluorescent molecule (Cy3) on the 5’ end and biotin on the 3’ end. ZF repressor templates were generated by PCR from gBlock gene fragments (IDT) that included the λ*P_R_* promoter. Global primers labeled with Cy5 at the 5’ end and biotin at the 3’ end were used to synthesize the ZF templates. To amplify ZF gene templates with cognate or non-cognate ligands a separate 3’ primer was used to incorporate the ligand. However, to produce ZF - PDZ or LZ gene templates, an assembly PCR was performed to link the sequence encoding the ZF with that encoding the PDZ or LZ domain. The two-finger homodimer *AA – GCN* was amplified from a gBlock gene fragment. All linear DNA templates contained 5’ protection sequences of 250 bp (promoters) or 130 bp (zinc fingers); this slows degradation of DNA in the cell-free extract (Sun et al. 2014). Complete sequences are given in Tables S3–S6. Given that each linear template is tagged by a fluorescent molecule at the 5’ end and a biotin molecule at the 3’ end we can assume that all DNA which is immobilized at the surface of a given unit cell and we visualize via fluorescence is therefore full length.

### Setting up high-throughput cell-free experiments

The concentration of linear template DNA was quantified by absorbance (NanoDrop, ThermoFisher) and PCR products were directly added to a 384 microwell plate along with short biotinylated ssDNA oligos for spotting. For characterization of the λ*P_R_* promoter library, a single linear template was spotted. Simple repression assays involved spotting both a ZF template along with the deGFP target template and dose response experiments for cooperative ZFs required spotting up to 5 linear templates per unit cell. Taking the PDZ-ligand heterodimer case as an example, both *ZF_BCB_ – PDZ* and *ZF_CBD_ – PDZ* as well as *ZF_ADD_ – L* and *ZF_CBD_ – NL* templates were combined in varying ratios, in addition to the deGFP template. All ZF templates and the target template were spotted for the NAND logic gate experiments, however for the AND and OR gates the first ZF template was pre-expressed in a separate reaction while all other down-stream components of the logic gate were spotted. Linear DNA templates were spotted with a QArray2 (Genetix) microarrayer using an MP3 pin (Arrayit) onto epoxy coated glass slides. The DNA was diluted in 4% BSA in MilliQ H_2_O to prevent the DNA from binding the epoxy functional groups and to aid visualization of the drops for alignment with the PDMS chip. Once the PDMS chip was aligned on top of the DNA microarray the chip was bonded to the glass by incubating the assembled device for 1 hour at 80 °C. Chips were then stored overnight at 45 °C and used the following day or up to 2-5 days later.

To prime the chip, control lines were filled with PBS and pressurized at up to 138 kPa. While isolating the spotted DNA with the neck valve closed, the lower half of the unit cell was patterned with BSA-biotin and neutravidin (Thermo Fisher) (Maerkl and Quake 2007). First, BSA-biotin (2 mg/mL) was flowed for 15 minutes, then neutravidin (1 mg/mL) for 15 minutes, after which the button was closed and BSA-biotin was flowed again for 10 minutes. Between each of these three steps PBS was flowed for 5 minutes to wash away any unbound molecules. The pressure applied to the flow lines during this process was ∼24 kPa. This surface chemistry resulted in a circular area coated with neutravidin, whereas the remaining surface of the unit cell was passivated with BSA-biotin. Afterwards the spotted DNA was solubilized with PBS by closing the outlet and opening the neck valve while PBS was flowed into the device. To avoid cross-contamination of DNA between unit cells the neck valve was closed and the lower half was washed with PBS. The sandwich valves were then closed and the neck was released to allow the DNA to diffuse into the lower half of the unit cell. After an incubation of 90 minutes the button was opened and the DNA was allowed to attach to the neutravidin coated area. The unbound DNA could then be washed away while the button was pressurized. The surface immobilized DNA was imaged by fluorescence microscopy. A 10 μL reaction volume of cell-free extract was prepared according to a previous protocol (Sun et al. 2013) and flowed into the device. For the AND and OR logic gates, the pre-synthesis reaction made up 25% of the fresh cell-free reaction mixture. The chip was separated into four parts using a multiplexer to test all logic gate inputs on a single chip. All unit cells were isolated from one another by the sandwich valves before the button was released and the chip was incubated at 29 °C. The production of deGFP in each unit cell was monitored over time on an automated fluorescence microscope (Nikon). The pneumatic valves, microscope, and camera were controlled by a custom LabVIEW program throughout the experiment.

### MITOMI measurements

DNA targets were either amplified with PCR or Klenow reactions and diluted over a range of 2 nM to 1 μM. The DNA was then spotted and the chip was aligned in the same way as described in the previous section. The same surface chemistry was also performed, followed by an additional two steps. First, a biotinylated anti-His antibody (Qiagen, 15 nM) was flowed for 20 minutes, enabling its binding to the neutravidin coated region. Second, a 30 μL cell-free reaction, in which a His-tagged ZF was expressed, was flowed for 25 minutes, immobilizing the ZF at the surface of each unit cell. The DNA targets were then solubilized and allowed to diffuse into the lower half of the unit cell where their binding to the ZF could be quantified via fluorescence microscopy. All image analysis was done using a custom MatLab script that either calculated the mean of the fluorescence signal at the button area or in the solution of the unit cell. *K_d_*s were determined by fitting the data with a single binding site model

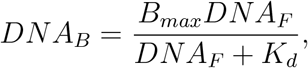

where *B_max_* is the maximum specific binding, *DNA_F_* is the free DNA in solution, *DNA_B_* is the DNA bound to protein at the surface and *K_d_* is the dissociation constant. Absolute *K_d_* values in molar units were subsequently determined according to a calibration made with known concentrations of Cy5-tagged DNA on-chip.

### Thermodynamic models for repression

Following (Bintu et al. 2005) we make the assumption that gene expression is proportional to RNAP occupancy, and that RNAP binding to promoters is at thermodynamic equilibrium. This allows the final protein expression level to be written as a function of the equilibrium occupancy of the promoters by RNAP, which itself is a function of the available microstates. Thus the model can be constructed by enumerating all possible binding states of RNAP and any transcription factor it interacts with. For concentrations of nonspecific sites and RNAPs given by *N_NS_* and *P*, respectively, a third assumption *P* ≪ *N_NS_* allows the occupancy to be written in the following simple form,

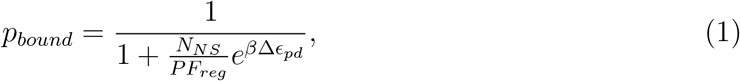

where the effects of the microstate distribution are subsumed into a regulation function *F_reg_*. Δ*є_pd_* is the energy difference between RNAP binding to specific versus nonspecific sites. All energies are given in units of *k_B_T*, and *β* = 1/*k_B_T*.

Typically repression is modeled using a purely competitive model where RNAP and the repressor competes for exclusive binding on the promoter. Motivated by our experimental observations, we extend this standard formulation of repression by enumerating four possible states for repressor binding, with the following energies:

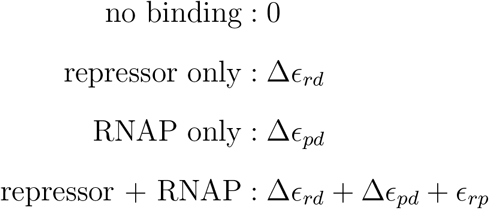

In our model, the repressor and RNAP can thus both bind at the same time, and interact with an energy *є_rp_*. As *є_rp_* becomes large and positive, the repression tends to that of competitive inhibition, where either species excludes the other from binding. This formulation allows for a continuous transition between competitive and noncompetitive mechanisms of inhibition: as repression becomes noncompetitive, simultaneous binding is possible and thus the promoter exhibits a non-zero leak at full repression.

We can then write down the regulation functions. For a single repressor of concentration *R* we have

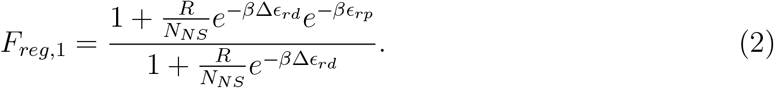

For repressors binding to two sites, we have

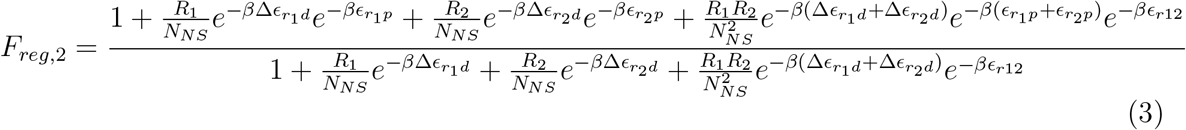

where the two repressors can interact with an energy *є*_*r*12_. Positive values of *є*_*r*12_ result in positive cooperativity, where the binding of one repressor facilitates the binding of the other.

We make two transformations to Equation 3. First, in dose response experiments, we assume that repressor concentration *R* is proportional to repressor DNA concentration *R* = *Bd_R_*, as supported by on-chip DNA titration measurements (Figure 1). This allows us to convert repressor concentrations to DNA concentrations, which is the experimentally varying quantity. Second, we simplify all our expressions by defining an effective dissociation constant

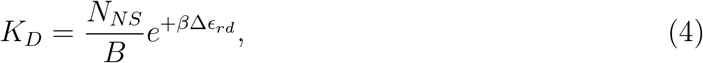

which gives the DNA, rather than the repressor concentration for half-maximum occupancy. The effective *K_D_*s in the model are related to standard physical dissociation constants *K_d_* = *e*^Δ*G*^Ө^/*k_B_T*^ by a multiplicative factor. These transformations result in the simplified equations

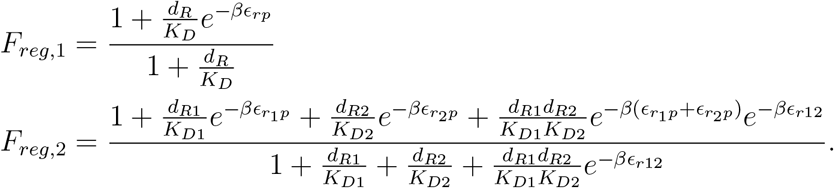

Finally, the protein level is given by a direct proportionality with the occupancy *y* = *Ap_bound_*, and fold repressions are given by ratios of protein levels. Thus,

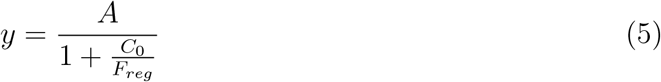

with

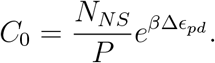

For binding site tuning experiments, the independent variable is an experimentally measured *K_d_*, in units of nM. The regulation functions become

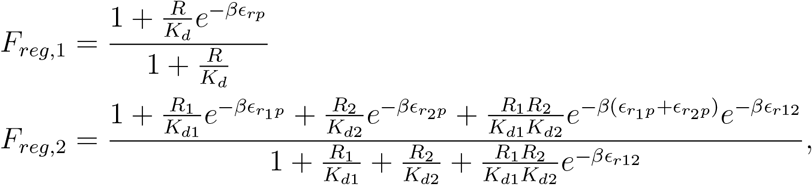

where *R* is now an effective repressor concentration, in units of nM.

To extend the model to take into account the distance-dependent effects, as well as the dependence of cooperativity on the helical positioning of the ZF on the DNA backbone, we posit the phenomenological description

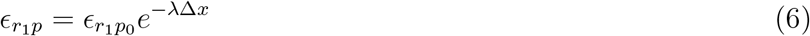

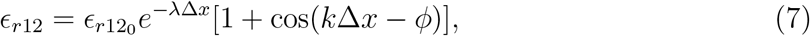

where λ is a distance decay constant, Δ*x* the spacing between the two ZF binding sites, *k* the wavenumber corresponding to the DNA helical pitch of 10.5 bp/turn, and *ϕ* a phase shift. The model was parameterized using values obtained from dose response measurements before fitting to the helical data. Finally, to take into account varying experimental conditions between those measurements and the helical effect experiments, a third parameter *R*_0_ was introduced as a multiplicative factor on DNA concentrations, 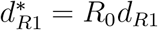 and 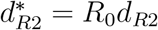. This parameterizes global changes such as a different total DNA concentration and protein production rates between the dose response and helical experiments.

### Markov chain Monte Carlo inference of model parameters

Model fits to experimental data are carried out using Markov chain Monte Carlo (MCMC) sampling, using the python package `emcee` (Foreman-Mackey et al. 2013) which is an implementation of Goodman and Weare’s affine-invariant ensemble sampling method (Goodman and Weare 2010). We first found a maximum likelihood estimate (MLE) of parameters using the BFGS algorithm from the `scipy.optimize` package. We define our negative log-likelihood function as

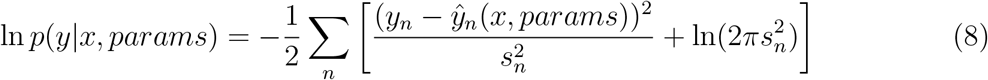

where *ŷ_n_*(*x, params*) is the model prediction and *y_n_* the experimental data with errors *s_n_*. In order to combine results from different experiments (for example, separate dose response curves), we added together the log-likelihood functions without normalization.

Uninformative, broad normal priors were used, centered on the MLE parameter values. 50 Markov chains were initialized in a tight ball around these MLE values, and allowed to run for 10,000 iterations. The first 5,000 points were considered as part of the burn-in period and discarded; equilibriation of the Markov chains was verified by inspection of the traces. Sampling the equilibrated Markov chains returns the posterior probability distributions of the parameters, which can be used to generate an ensemble of potential models (and hence, a distribution of fits as shown in Figure 5). The posterior distributions, as well as pairwise distributions of parameters are shown in Figure S9.

## SUPPLEMENTAL ITEMS

**Table S1:** Quantitative measures of cooperative interactions.

| Design: PDZ | Sensitivity | Hill coefficient |
| --- | --- | --- |
| BCB+ADD cooperative | 0.592±0.007 | 1.3±0.2 |
| BCB+ADD non-cooperative | 0.312±0.003 | 1.0±0.2 |
| BCB | 0.165±0.001 | 1.1±0.4 |
| ADD | 0.183±0.001 | 1.1±0.4 |

| Design: LZ | Sensitivity | Hill coefficient |
| --- | --- | --- |
| BCB+ADD cooperative | 0.627±0.009 | 1.3±0.2 |
| BCB+ADD non-cooperative | 0.422±0.002 | 1.1±0.3 |
| BCB | 0.166±0.001 | 1.2±0.4 |
| ADD | 0.182±0.001 | 1.3±0.5 |

**Table S2:** Inference of parameter values. The posterior probability distribution of each parameter was quantified by reporting the value for the 50^th^ percentile (large numbers) as well as uncertainty bounds given by the 16^th^ and 84^th^ percentiles (small numbers).

Dose response fits
| Parameters: PDZ | Value | Units |
| --- | --- | --- |
| A | $+5.76e + 03^{+8.02e+02}_{-7.19e+02}$ | RFU |
| C<sub>0</sub> | $+6.62e - 01^{+2.28e-01}_{-1.99e-01}$ | |
| K<sub>1</sub> | $+1.66e - 01^{+7.12e-02}_{-5.09e-02}$ | |
| K<sub>2</sub> | $+8.24e - 03^{+4.67e-03}_{-3.01e-03}$ | |
| ε<sub>r1p</sub> | $+1.43e + 00^{+1.81e-01}_{-1.78e-01}$ | k<sub>B</sub>T |
| ε<sub>r2p</sub> | $+1.32e + 00^{+1.88e-01}_{-1.57e-01}$ | k<sub>B</sub>T |
| ε<sub>r12</sub> | $-3.46e + 00^{+4.36e-01}_{-4.66e-01}$ | k<sub>B</sub>T |

| Parameters: LZ | Value | Units |
| --- | --- | --- |
| A | $+6.26e + 03^{+7.54e+02}_{-6.95e+02}$ | RFU |
| C<sub>0</sub> | $+8.84e - 01^{+2.18e-01}_{-1.95e-01}$ | |
| K<sub>1</sub> | $+6.86e - 02^{+2.50e-02}_{-2.02e-02}$ | |
| K<sub>2</sub> | $+4.27e - 02^{+1.92e-02}_{-1.45e-02}$ | |
| ε<sub>r1p</sub> | $+1.28e + 00^{+1.86e-01}_{-1.74e-01}$ | k<sub>B</sub>T |
| ε<sub>r2p</sub> | $+1.19e + 00^{+1.77e-01}_{-1.54e-01}$ | k<sub>B</sub>T |
| ε<sub>r12</sub> | $-3.05e + 00^{+3.92e-01}_{-3.99e-01}$ | k<sub>B</sub>T |

| Parameters: AA-LZ | Value | Units |
| --- | --- | --- |
| A | $+5.13e + 03^{+8.87e+02}_{-8.27e+02}$ | RFU |
| C<sub>0</sub> | $+5.70e - 01^{+2.77e-01}_{-2.48e-01}$ | |
| K | $+8.42e - 01^{+4.79e-01}_{-3.16e-01}$ | |
| ε<sub>rp</sub> | $+1.44e + 00^{+1.98e-01}_{-1.35e-01}$ | k<sub>B</sub>T |
| ε<sub>r12</sub> | $-9.48e + 00^{+7.85e-01}_{-8.39e-01}$ | k<sub>B</sub>T |

| Parameters: Helical | Value | Units |
| --- | --- | --- |
| ln(λ) | $-3.41e + 03^{+1.18e-01}_{-1.16e-01}$ | |
| φ | $-1.92e + 00^{+1.21e-01}_{-1.14e-01}$ | |
| ln(R<sub>0</sub>) | $-2.24e + 00^{+1.78e-01}_{-1.72e-01}$ | |

Binding site tuning fits
| Parameters: AAA | Value | Units |
| --- | --- | --- |
| C<sub>0</sub> | $+1.55e - 02^{+1.93e-03}_{-1.57e-03}$ | |
| R | $+4.94e + 02^{+5.04e+01}_{-5.09e+01}$ | nM |
| ε<sub>rp</sub> | $+1.13e + 01^{+2.92e+00}_{-1.94e+00}$ | k<sub>B</sub>T |

| Parameters: BCB | Value | Units |
| --- | --- | --- |
| C<sub>0</sub> | $+1.35e - 02^{+1.62e-03}_{-1.33e-03}$ | |
| R | $+4.93e + 02^{+5.03e+01}_{-5.05e+01}$ | nM |
| ε<sub>rp</sub> | $+1.20e + 01^{+2.78e+00}_{-1.85e+00}$ | k<sub>B</sub>T |

| Parameters: AA-LZ | Value | Units |
| --- | --- | --- |
| $C_0$ | $+2.82e - 03^{+1.64e-03}_{-1.15e-03}$ | |
| $R$ | $+9.78e + 01^{+1.00e+01}_{-1.01e+01}$ | nM |
| $\epsilon_{rp}$ | $+1.12e + 01^{+7.33e-01}_{-6.70e-01}$ | $k_B T$ |
| $\epsilon_{r12}$ | $-4.12e + 00^{+4.72e-01}_{-5.37e-01}$ | $k_B T$ |

| Parameters: BCB/ADD | Value | Units |
| --- | --- | --- |
| $C_0$ | $+1.64e - 02^{+2.38e-02}_{-1.10e-02}$ | |
| $R_1$ | $+9.08e + 01^{+9.06e+00}_{-8.80e+00}$ | nM |
| $R_2$ | $+5.00e + 01^{+4.81e+00}_{-5.09e+00}$ | nM |
| $\epsilon_{r1p}$ | $+1.04e + 01^{+9.03e-01}_{-8.68e-01}$ | $k_B T$ |
| $\epsilon_{r2p}$ | $+1.06e + 00^{+9.13e-01}_{-7.10e-01}$ | $k_B T$ |
| $\epsilon_{r12}$ | $-1.45e + 00^{+5.34e-01}_{-5.94e-01}$ | $k_B T$ |

**Table S3:**
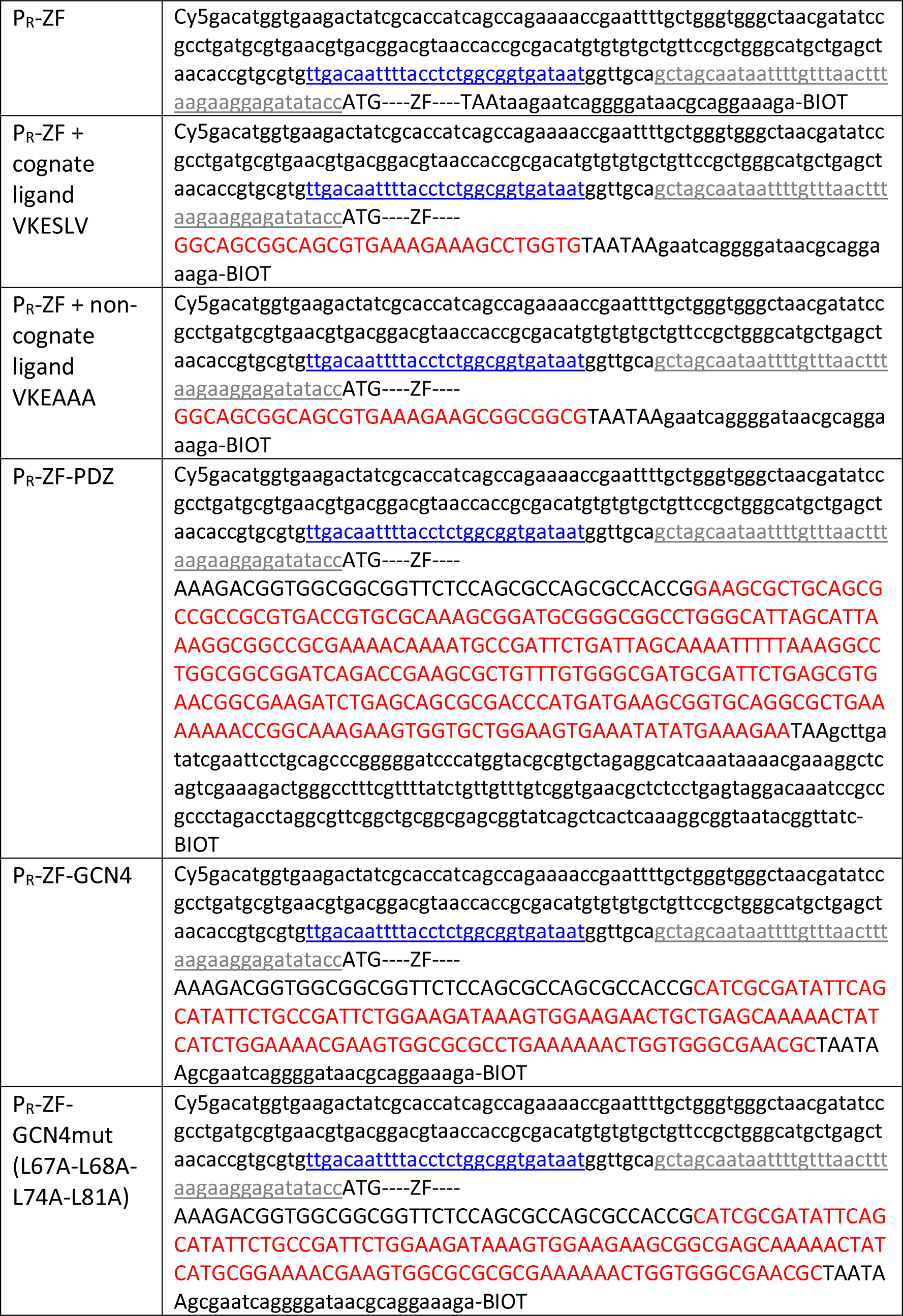
ZF expression templates. 3-finger ZFs were expressed from lambda P_R_ promoters (blue, underlined) with conserved 5’UTR (grey, underlined), and could be modified to contain interaction domains and ligands (red).

**Table S4:**
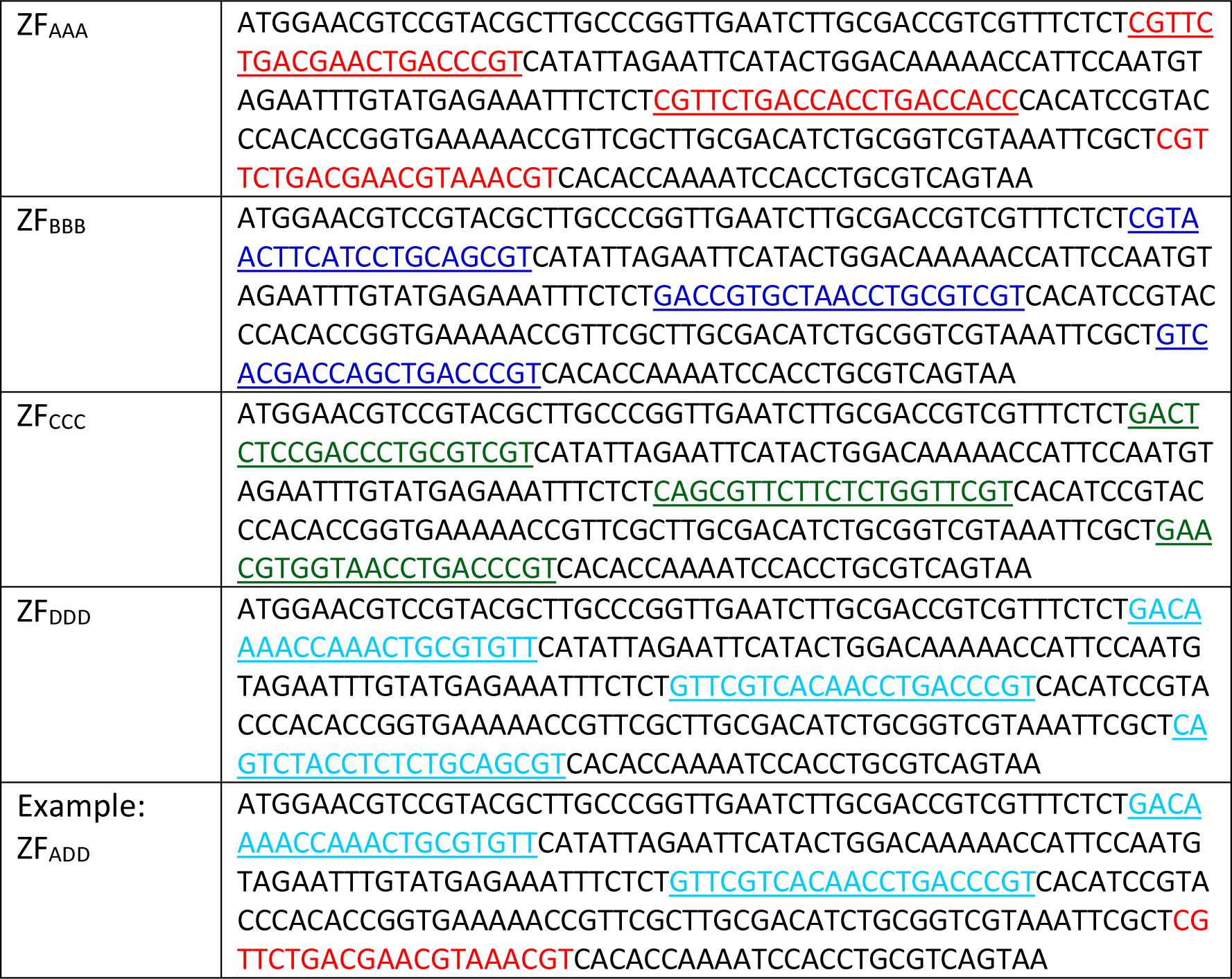
Three-finger ZF coding sequences. Variable recognition helices for each finger (1-3) are highlighted. ZFs bind in the 3’-5’ direction by convention, and thus to construct, for example, ZF_ADD_ the fingers must be arranged in reverse order F1 (D)-F2 (D)-F3 (A).

**Table S5:**
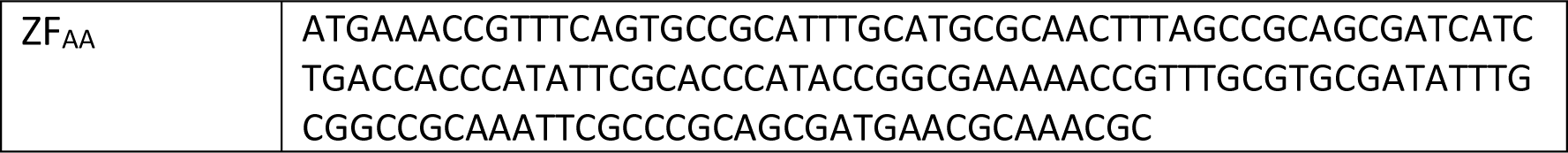
Two-finger ZF coding sequence.

**Table S6:**
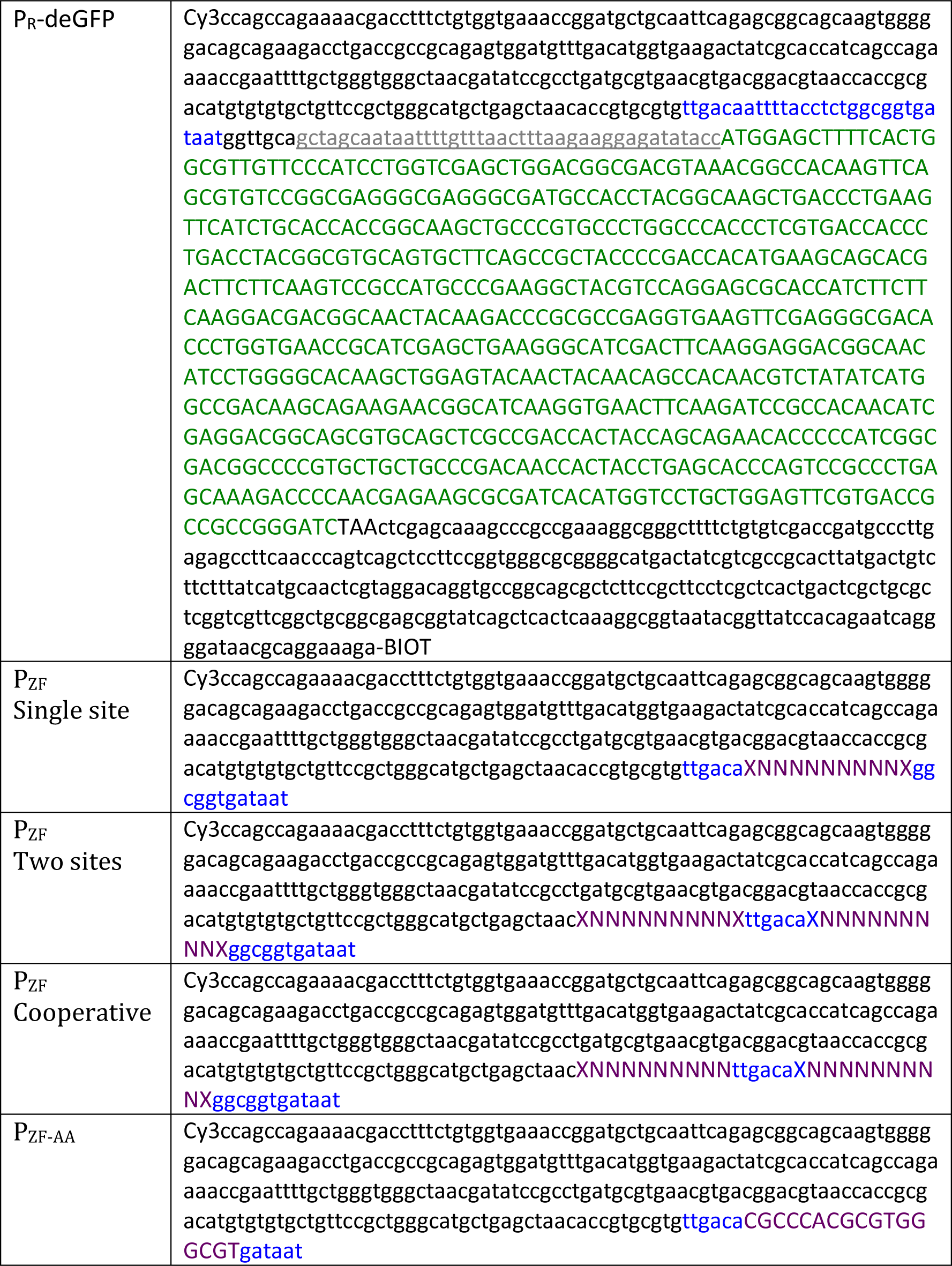
Reporter and target promoter design. deGFP was used as a reporter. The upstream regulatory region consisted of variations of the original lambda P_R_ promoter containing 9-bp ZF binding sites (N) and their flanking bases (X).

**Table S7:** Three-finger ZF target sequences. Each ZF binds a 9bp target; for each target we conserve single flanking bases on either side where possible.

|  |  |
| --- | --- |
| Target AAA | 5'-a GCG TGG GCG t-3' |
| Target BBB | 5'-t GAG GAC GTG t-3' |
| Target CCC | 5'-a GAT GTA GCC t-3' |
| Target DDD | 5'-t GTA GAT GGA g-3' |
| Example:<br>target ADD | 5'-a GCG GAT GGA g-3' |

**Table S8:**
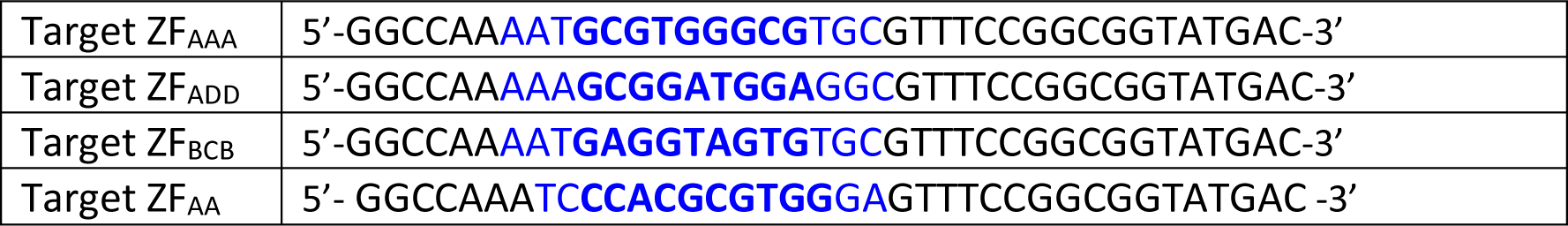
MITOMI PWM targets. The bases which were mutated are colored blue and the bold bases represent the core binding sequence.

